# Specificity and tunability of efflux pumps: a new role for the proton gradient?

**DOI:** 10.1101/2024.06.11.598359

**Authors:** Matthew Gerry, Duncan Kirby, Boian S. Alexandrov, Dvira Segal, Anton Zilman

## Abstract

Bacterial efflux pumps that transport antibacterial drugs out of the bacterial cells have broad specificity, commonly leading to broad spectrum resistance and limiting treatment strategies for infections. It remains unclear how efflux pumps can maintain this broad spectrum specificity to diverse drug molecules while limiting the efflux of other cytoplasmic content. We investigate the origins of this broad specificity using theoretical models informed by the experimentally determined structural and kinetic properties of efflux pumps. We develop a set of mathematical models describing operation of efflux pumps as a discrete cyclic stochastic process across a network of states characterizing pump conformations and the presence/absence of bound ligands and protons. We find that the pump specificity is determined not solely by the drug affinity to the pump–as is commonly assumed–but it is also directly affected by the periplasmic pH and the transmembrane potential. Therefore, the pump effectiveness in removing a particular drug molecule from the cell can be tuned by modifying the proton concentration gradient and the voltage drop across the membrane. Furthermore, we find that while both the proton concentration gradient across the membrane and the transmembrane potential contribute to the thermodynamic force driving the pump, their effects on the efflux enter not strictly in a combined proton motive force, but rather they have two distinguishable effects on the overall throughput. These results potentially explain the broad specificity of efflux pumps and suggest ways to overcome bacterial resistance, while highlighting unexpected effects of thermodynamic driving forces out of equilibrium.

## I. INTRODUCTION

In recent decades, bacterial antibiotic resistance has emerged as a major and pervasive threat to public health and, consequently, it has become a critical subject of study of high biomedical significance [1–4]. Bacterial drug resistance is a complex, dynamic response to the evolutionary pressure generated by the widespread use of antibiotics in modern society [3, 4]. In other words, bacteria rapidly evolve, predominantly via horizontal gene transfer [5, 6], to diversify and optimize their defense mechanisms and to neutralize new antimicrobial agents [7]. Multidrug transmembrane active transporters, commonly known as bacterial efflux pumps, play a central role among these defense mechanisms. These pumps are effective against a wide range of different drugs, while at the same time exhibiting the specificity needed to preferentially extrude drug molecules, rather than self molecules, from the cell [7–13].

In this study, we focus on minimal models for active transmembrane transporters, or efflux pumps, that are powered by proton concentration gradients across the inner membrane of the cell and can pump a variety of different drug molecules. In doing so, they confer multidrug resistance to bacterial cells. The models we consider are inspired loosely by various families of transporters known to be associated with multidrug resistance [11]. Resistance-Nodulation-Division (RND) efflux pumps comprise one such family, which includes the AcrAB-TolC pump found in *Escherichia coli* and the similarly structured MexAB-OprM pump found in *Pseudomonas aeruginosa*. RND pumps are elaborate protein complexes with a trimeric structure giving rise to a “rotary” pumping mechanism [14–19]. Another relevant class of pumps consists of symporters and antiporters of the Major Facilitator Superfamily (MFS), which instead exhibit an alternating-access mechanism [20]. To study the general properties of efflux drug specificity, efficiency, and tunability common across different transporter classes, we subsume many of these molecular features into simpler, more abstract models whose kinetics can nevertheless serve as a good approximation to those of these more realistic pumps [21]. The conclusions of the study of these minimal models also bear on the understanding of the operation of other antiporter pumps, involved in, for example, maintaining proton and sodium ion concentration gradients across membranes [22–24].

Schematics of pump operation are depicted in Fig. 1. The pump spans the inner and outer membrane of the cell, with two functionally and spatially distinct components. The outer-membrane component is a passive exit conduit responsible for the diffusion of the antimicrobial agents into the extracellular space [25]. The inner-membrane component is responsible for the active pumping of the antimicrobial agents out of the cytoplasm via coupling to the flux of the protons. This flux is, in turn, driven by electric and chemical potential gradients across the membrane, which are present due to the relatively high concentration of protons within the periplasm, or intermembrane space. In light of this proton flux, it is implicit that other processes in the cell must dissipate energy in order to maintain these nonequilibrium gradients. This amounts to a thermodynamic cost the cell must pay to ensure that the pump preferentially delivers drug molecules from the cytoplasm towards the exit duct, regardless of the direction of the concentration gradient of the drug molecules themselves.

**FIG. 1.**
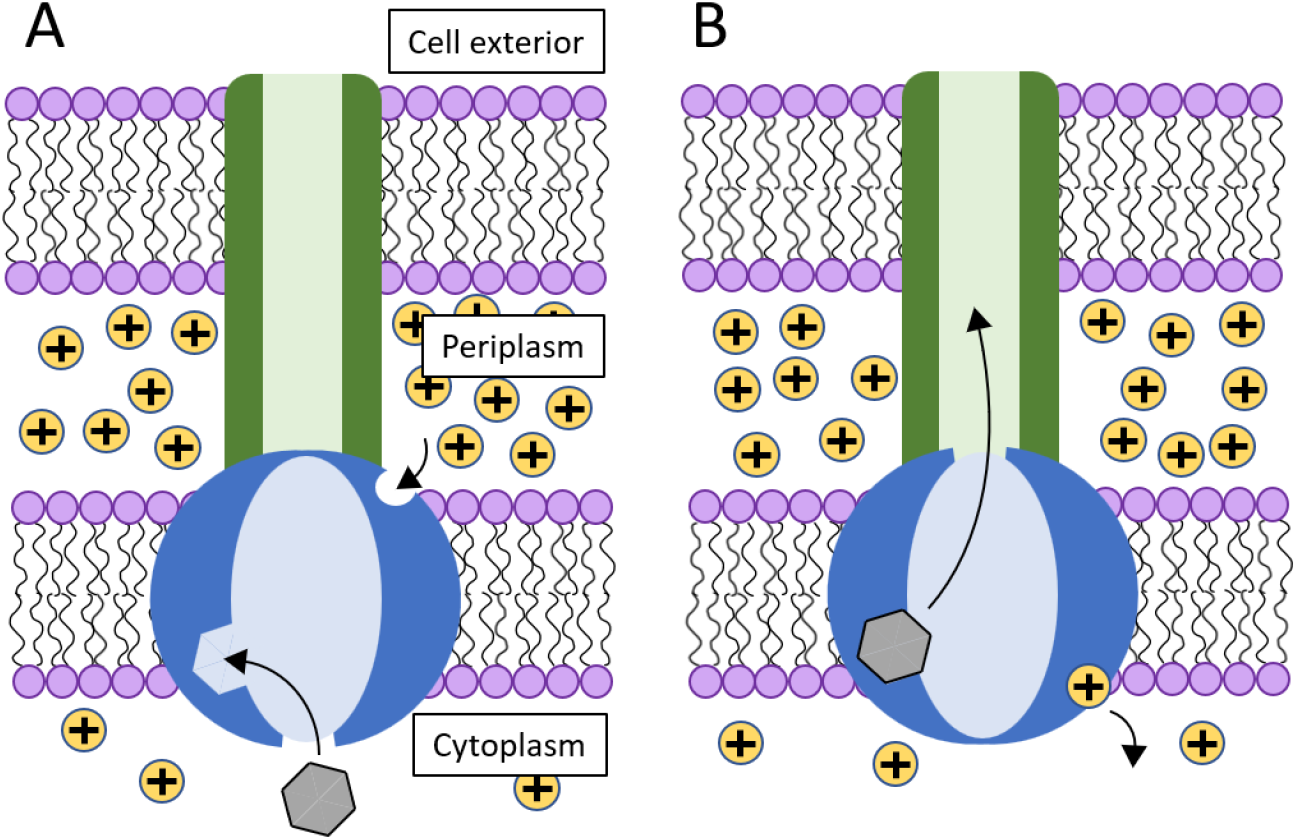
Schematic depiction of the proton antiporter pump considered in this work in each of its two conformational states. The active component of the pump is in blue, coupled to the passive efflux channel shown in green. Arrows indicate the typical motion of ligands throughout the pumping process. **A**. A drug molecule (gray) binds from the cytoplasm to a binding site on the pump interior. A proton binds to its respective binding site, initially facing the proton-rich periplasm. **B**. Upon proton translocation toward the cytoplasm, the channel opens to the passive transporter (green) which allows the drug to exit the cell.

We present two mathematical models of progressing complexity that describe the operation of the active inner-membrane transporter as a Markov jump process on a network of states, with transitions corresponding to the binding and unbinding of ligands and changes in pump conformation. The use of such models demands the identification of a finite set of coarse-grained states approximating the operation of the system, which, in reality, evolves continuously. These states are typically taken to represent the most stable and long-lived molecular conformations as determined through structural measurements. Numerous studies have employed models of this kind to describe the operation of various protein complexes, including ATP-powered pumps [26–28], molecular motors [29–31], kinetic proofreading receptors [32, 33], and DNA polymerases [34, 35].

Our aim is to shed light on how an efflux pump can effectively transport a wide a range of drug molecules with differing properties while simultaneously exhibiting the specificity necessary to avoid pumping out the molecules needed for the normal functioning of the cell. We show that, as expected, the periplasmic pH, the membrane potential, and the strength of proton and drug binding to the transporter all play a role in modulating the flux throughput of the pump. Notably, unlike the expectations based on the Onsager theory close to equilibrium [36], the membrane potential and the pH modulate the pump throughput not strictly through their combination in the electrochemical potential of protons (broadly known as the proton motive force), but with each having their own distinct effect.

Furthermore, tuning membrane potential and periplasmic pH parameters impacts not only the overall efflux rate, but also pump specificity, by shifting the range of drug binding affinities at which the pump throughput is maximized. This suggests a mechanism by which the bacterial population can tailor its response to drugs used in treatment, and thus may explain the hitherto puzzling broad specificity of the multi-drug efflux pumps. In light of these findings, we discuss the functional cycle of the transporter and the coupling of its specificity and efficiency with the proton gradient, and identify possible considerations towards designing more effective drug design protocols [7].

This paper is structured as follows. In Section IIII A we introduce and describe a minimal three-state kinetic model of efflux pumps. In Section IIII B we discuss the different factors affecting the overall efflux as predicted by the model, in particular the nontrivial dependence of the efflux on the proton concentrations in the periplasm and cytoplasm, and on the membrane electric potential. In Section IIII C we demonstrate how changes to the periplasmic pH can tune the range of drug binding affinities at which the pump is most effective, suggesting a mechanism by which efflux pump specificity can be tuned in response to drugs used in treatment. Section IIII D introduces a richer kinetic model which exemplifies the same effects, broadening the class of system to which this analysis may apply. Section III includes a discussion of how our work highlights nontrivial effects seen in cellular processes taking place far from equilibrium, while fitting into the context of the existing understanding of multidrug resistance, and informing the development of potential future treatments for bacterial infections.

## II. RESULTS

### A. Three-state kinetic model

We consider a simple model for a proton antiporter pump as depicted in Fig. 1, with one binding site for the drug molecule initially facing the interior of the cell and one binding site for protons facing the periplasm. An analogous pump with multiple proton binding sites could be considered as a straightforward generalization. After the sequential binding of the drug molecule and the proton [19], a conformational change occurs in the pump as the proton is pulled by the electric potential gradient across the inner membrane from the periplasmic side towards the cytpolasmic side, coupled with the release of the drug into the cytoplasm. This exposes the ligands to their respective destinations: the drug molecule is able to unbind and be released to a passive channel that carries it out of the cell, and the protons are released to the cytoplasm. Note that in the figure, the source/destination of each ligand is denoted in parentheses.

To model the operation of this pump, we consider a three-state kinetic scheme, as shown in Fig. 2A. The associated process characterizing one completion of the cycle depicted consists of three steps. As required for microscopic thermodynamic consistency, each of these steps can occur in either direction.

**FIG. 2.**
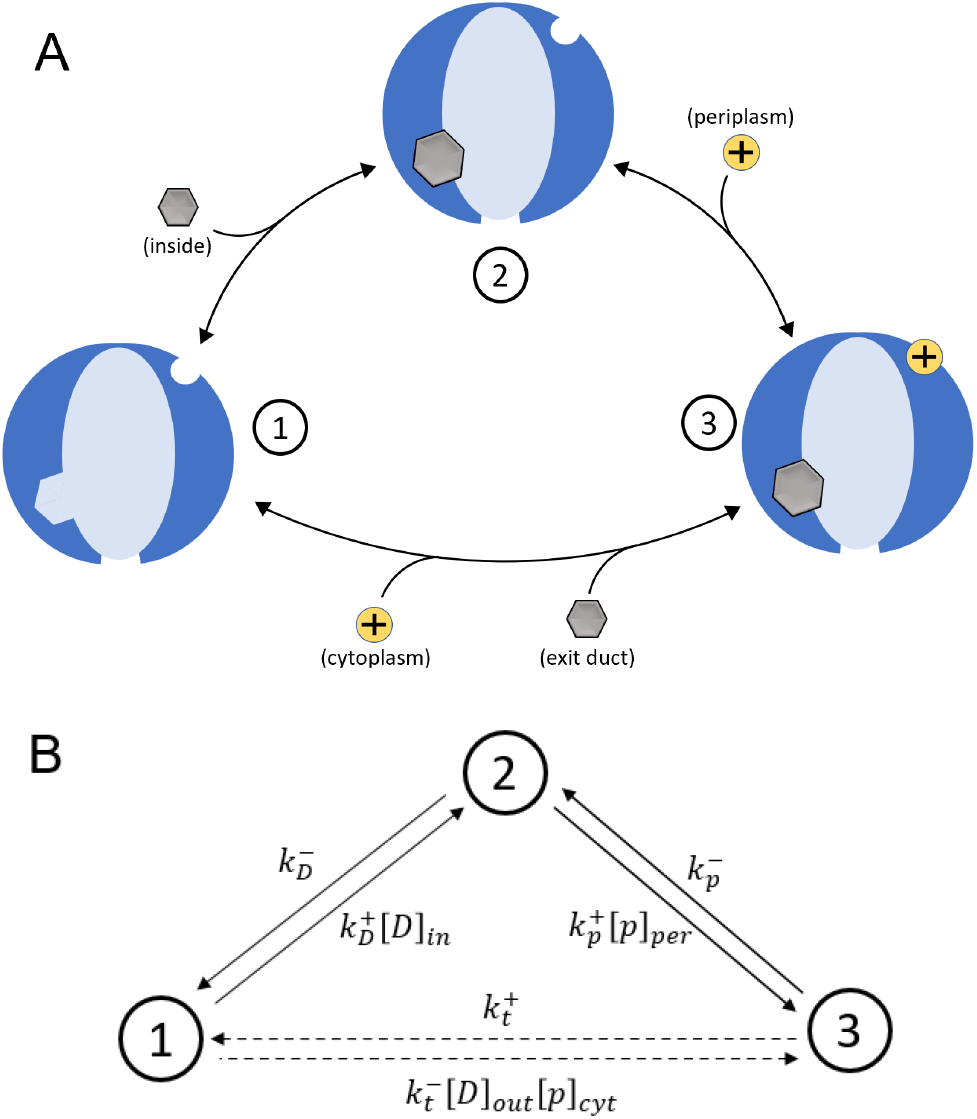
**A**. Three-state kinetic model of the efflux pump, with reversible transitions represented by black arrows. Drug molecules and protons are shown where they bind to (unbind from) the pump, labelled with the relevant reservoir in each case. The forward process proceeds in the clockwise direction. **B**. The model represented in terms of directed transitions of a network of states, with associated transition rates shown. The mathematical formulation of the model is presented in Equation (1) and the kinetic parameters are defined in Equations (2)-(5).

In state 1, neither a drug molecule nor any protons are bound. Then, in a transition to state 2 a drug molecule binds to its respective binding site. We consider a pump with drug binding from the cytoplasm, as depicted in Fig. 1, but we note that a proton antiporter pump like this may have multiple different drug binding sites associated with different drug molecular mass ranges. Drug binding may, in actuality, occur from either the cytoplasm or the periplasm [37–39]. Subsequently, a proton binds to its binding site taking the pump to state 3. The third and final step, which takes the pump back to its initial state, consists of four key sub-processes. Firstly, a conformational change occurs, closing the pump off to the interior of the cell and opening it to the exit duct, while the protons are simultaneously transported across the inner membrane towards the cytoplasm. Then the drug unbinds and is released to the exit duct. The protons unbind and exit into the cytoplasm. Finally, in the absence of bound ligands, the pump returns to its original conformation, with the drug binding site exposed to the cell interior. Despite using an alternating-access mechanism rather than a rotary mechanism, this sequence of transitions between of coarse-grained states is motivated by experiments on pumps of the RND family [19]. However, we note that the model itself is meant to be a simplified idealized representation of a broad class of transporters.

The simplicity of this model allows us to obtain analytic results pertaining to the behaviour of efflux pumps. However, this simplicity is also the source of this model’s limitations: namely, the strong assumption that multiple events always occur in tandem comprising the final 3 → 1 transition. Relaxation of some of these constraints, and corresponding expansion of the parameter space, leads to a model that, while less mathematically tractable, may describe the operation of certain efflux pumps with greater accuracy. We explore this notion in Sec. IIII D.

The system is described by the probabilities, *P*_1_, *P*_2_, and *P*_3_, of the transporter to be in each of the three states, which are governed by the corresponding Master equation that describes the transitions between states. Conceptually similar models have been widely and successfully used in the past to describe active transporters [24, 40–45]. Accordingly,

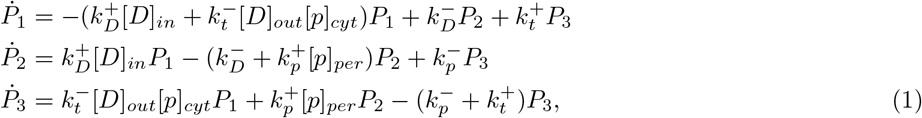

where 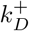 and 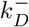 are forward and reverse rate constants for the drug binding step, 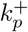 and 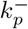 are those for the proton binding step, and 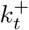 and 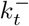 are those for the final, combined step. [*D*]_*cyt*_ and [*p*]_*per*_ are the cytoplasmic concentration of the drug and periplasmic concentration of the protons, respectively (−log_10_[*p*]_*per*_ is the periplasmic pH). The master equation is represented graphically in Fig. 2B, where each transition is labelled with its respective rate. Rates in Eqs. (1) for transitions that involve drug or proton binding are proportional to the relevant ligand concentration, following a law of mass action [46]. We take the rate at which the pump transports drug molecules and protons to be slow in comparison to other translocation processes happening in the cell, allowing concentrations to be assumed constant.

[*D*]_*out*_ and [*p*]_*cyt*_ are the extracellular drug concentration and cytoplasmic proton concentration. These factor into the transition rates for the backwards operation of the pump, which is rare under typical conditions, due to the electric potential gradient playing the strongest role in determining the directionality of the pump operation.

The transition rates are expressed in terms of parameters characterizing the system. For the drug binding step, the ratio of the binding and unbinding rates is equal to the dissociation constant describing the drug interaction with the pump:

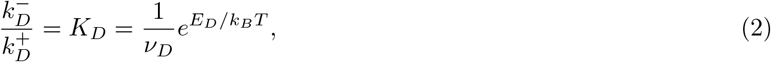

where *k*_*B*_ is the Boltzmann constant, *T* is the temperature of the surroundings, *E*_*D*_ *<* 0 is the drug binding energy to the pump, and *ν*_*D*_ is a constant related to the volume and the shape of the interacting moieties.

Similarly, for the interaction of the protons with the protonatable groups in the inner-membrane domain:

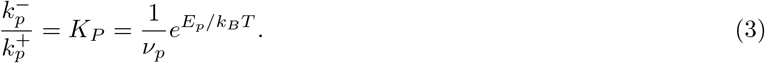

The constant *K*_*p*_ is linked to the protons entering the transmembrane domain and to their transient weak interaction with the protonatable residues therein.

We take the drug binding rate to be 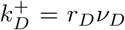, where *ν*_*D*_ is an interaction volume and 1*/r*_*D*_ is a characteristic timescale for a molecule to bind once in proximity to its binding site (as set by the interaction volume). This describes the case where drug-pump association is dominated by diffusion. *r*_*D*_ captures the effects of potential energy barriers between coarse-grained states [43], and the unbinding rate is 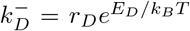, ensuring detailed balance relations are satisfied with respect to the drug concentration gradient [46].

Similarly for the protons, 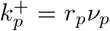 and 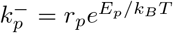, with *r*_*p*_ and *ν*_*p*_ defined analogously. While the physical interpretation of these quantities for proton binding is less clear, defining them provides the basis for exploring the entire parameter space of the kinetic model. We note that the forward and reverse rates for each step must satisfy *local* detailed balance relations, as set by the thermodynamic force relevant to the individual transition. This accounts, for instance, for the distinction between the factor 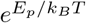 appearing in 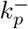 and the factor 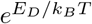 appearing in 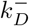. As a result, the entire system does not satisfy a global detailed balance relation, reflecting the fact that it operates away from equilibrium and gives rise to nonzero fluxes at steady state [47].

In the simplified model of Fig. 2, the final transition from state 3 to 1 comprises four separate events. However, in spite of its complexity, the rate of this transition can be compactly related to differences in the energies of states 1 and 3 as follows:

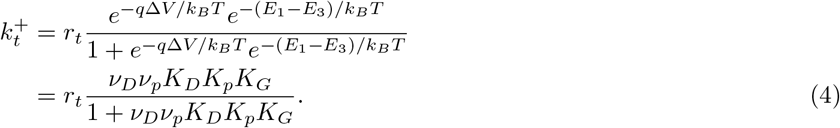

Since the system returns to its initial state after this step, it follows that *E*_1_ − *E*_3_ = −*E*_*D*_ − *E*_*p*_, accounting for the change in internal energy. We substitute these energy values for other parameters via Eqs. (2) and (3). −*q*Δ*V* is the change in the electrostatic energy of the protons upon translocation into the cytoplasm, and *K*_*G*_ = exp(−*q*Δ*V/k*_*B*_*T*) is the associated Boltzmann factor. Eq. (4) reflects the fact that the electric potential acts during the conformational change, forcing the proton from the periplasm to the cytoplasm. The form of the denominator ensures that the expected behaviour occurs in various limits (e.g. weak/strong binding, membrane potential).

The rate constant 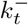 for reverse transitions 1 → 3 is then fixed by local detailed balance relations as [47, 48]

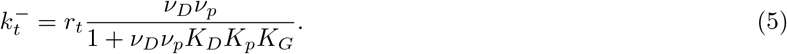

We note the presence of a free parameter *r*_*t*_ with units of inverse time, analogous to *r*_*D*_ and *r*_*p*_ but without interpretation as the characteristic rate for any single process. It is necessary to capture microscopic details not explicitly accounted for in the model while providing the richness to explore the parameter space as needed. This parameter is no longer needed for the richer model discussed in Sec. IIII D.

The rate *J* of drug efflux generated by the pump at steady state can be calculated as the net population flux from state 3 to state 1 in the kinetic scheme:

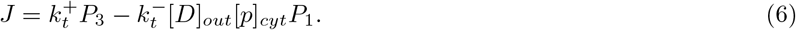

The steady-state populations *P*_1_ and *P*_3_ can be obtained from the system of equations (1) by setting each time derivative 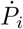 to zero.

### B. Pump throughput is determined by more than just the drug binding affinity to the pump and the proton motive force

Upon solving the system of equations (1) and evaluating Eq. (6), the efflux rate *J* can be written in terms of the model parameters as,

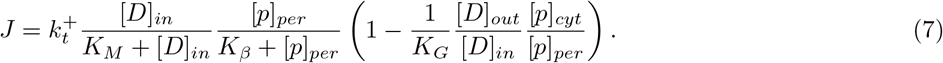

In the limit that *K*_*G*_ ≫ [*D*]_*out*_[*p*]_*cyt*_*/*[*D*]_*in*_[*p*]_*per*_, i.e., the *irreversible* limit, the factor in brackets tends to 1 and the efflux rate *J* described by Eq. (7) reproduces the experimentally observed Michaelis-Menten (MM)-like dependence of the efflux rate on the drug concentration [49, 50].

*K*_*β*_ and *K*_*M*_ may be understood to be a pair effective Michaelis-Menten constants. The former is given by

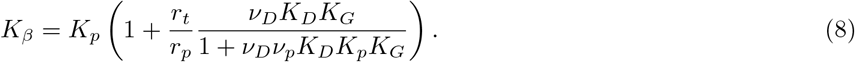

*K*_*M*_ may be viewed as the effective affinity with which the pump extrudes drug molecules. It is a complicated function of the characteristic rates, binding affinities, electric potential, outside drug concentration, and both proton concentrations (Supporting Information). Under experimentally relevant conditions (Supporting Information), it is approximated as

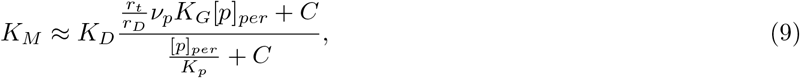

where 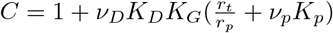.

Importantly, this effective affinity depends explicitly on the periplasmic concentration of the protons and the membrane potential, approaching the value of *K*_*D*_ only in the limit of very low [*p*]_*per*_ and otherwise varying dramatically with the other environmental parameters. This reflects that, in general, the drug affinity towards the pump is not the only determinant of its specificity.

The dependence of the efflux on the inside drug concentration and *K*_*D*_ is illustrated in Fig. 3A which shows normalized non-dimensional efflux 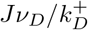, representing the efflux rate relative to the characteristic rate for a drug molecule in the interaction volume to diffuse to the binding site. The aforementioned Michaelis-Menten kinetics are demonstrated, with the efflux rate saturating as the drug concentration inside the cell increases, particularly since *K*_*G*_ tends to be large, setting the bracketed factor in Eq. (7) close to 1, corresponding to the irreversible limit of pump operation. Furthermore, the flux saturates to different levels for different *K*_*D*_’s indicating the ability of the pump to discriminate between different molecules independent of their concentration, known as “absolute discrimination” [51, 52].

**FIG. 3.**
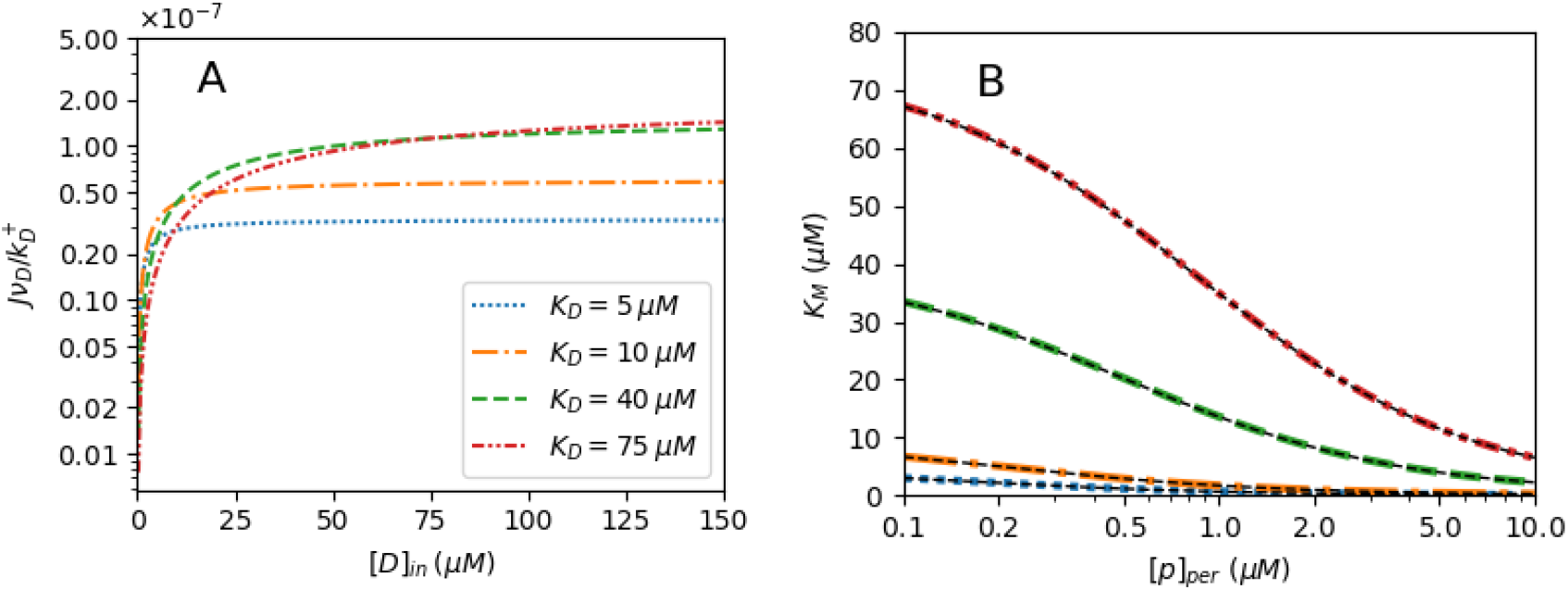
**A**. The dimensionless efflux, 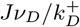, as a function of [*D*]_*in*_ for varying *K*_*M*_. The efflux increases monotonically with [*D*]_*in*_, as predicted by Eq. (7) and consistent with Michaelis-Menten kinetics. The value of *K*_*D*_ influences the drug concentration level at which the efflux rate begins to saturate, as well as the value to which it saturates. [*p*]_*per*_ = 1 *µM*. **B**. The Michaelis-Menten constant, *K*_*M*_, as a function of the periplasmic proton concentration. The narrow, black, dashed lines represent the simplified expression, Eq. (9), showing agreement with the exact values. It is approximated reasonably well by the drug dissociation constant, *K*_*D*_, only in the limit of low [*p*]_*per*_, and otherwise varies with variations in the periplasmic pH, membrane voltage, and characteristic rates. [*D*]_*in*_ = *K*_*D*_ = 10 *µM*. In both panels, [*D*]_*out*_ = 10 *µM* and [*p*]_*cyt*_ = 0.1 *µM*. The remaining parameter values are set as *T* = 295 *K, r*_*D*_ = 10^8^ *s*^−1^, *r*_*p*_ = 10^14^ *s*^−1^, *r*_*t*_ = 10^17^ *s*^−1^, *v*_*D*_ = 1 *M* ^−1^, *ν*_*p*_ = 10^−6^ *M* ^−1^, *K*_*p*_ = 0.1 *µM*, and *K*_*G*_ = 100 (Supporting Information).

Under the parameter values in the ranges discussed above, various simplifications (for example, *ν*_*D*_*ν*_*p*_*K*_*D*_*K*_*p*_*K*_*G*_ ≪ 1) lead to the derivation of Eq. (9). One can see by inspecting this expression that, in this regime, *K*_*M*_ is approximated reasonably well by the drug dissociation constant, *K*_*D*_, only in the limit of low periplasmic proton concentration. This behaviour is shown in Fig. 3B–*K*_*M*_ is seen to deviate from *K*_*D*_ as the proton concentration in the periplasm grows.

A few insights concerning the factors affecting the efflux rate can be drawn here from the functional form of *J*. Focusing first on the dependence of the efflux on drug concentrations, we note that the factor in the brackets recovers the expected linear dependence on the concentration gradient, Δ*µ*_*D*_ = *k*_*B*_*T* ln([*D*]_*out*_*/*[*D*]_*in*_), in the near-equilibrium regime [36]. However, [*D*]_*in*_ also appears in the factor describing the MM-like behaviour, here as part of the ratio [*D*]_*in*_*/K*_*M*_. The effective affinity *K*_*M*_, in turn, depends strongly on the membrane potential, as well as the proton concentration gradient via its direct dependence on [*p*]_*per*_, outside of the special case that *r*_*t*_*ν*_*p*_*K*_*p*_*K*_*G*_*/r*_*D*_ ≈ 1 (Supporting Information). We stress that this is a direct manifestation of the nonequilibrium nature of the pump’s operation, distinct from the electrochemical gradients simply amounting to the thermodynamic forces driving the efflux. The complicated behaviour of *K*_*M*_ stands in sharp contrast to any equilibrium situation, where the dissociation constant, *K*_*D*_, is the sole quantity relevant to characterizing the drug’s interaction with the pump.

Turning attention to the effects of the proton concentrations, we identify the electrochemical potential, or “proton motive force” (p.m.f.) as the quantity most often assumed relevant to the operation of proton-powered molecular motors,

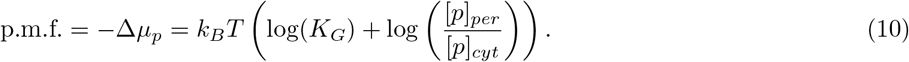

As for the drug concentration gradient, the membrane potential and proton concentrations do appear in this form in the bracketed factor in Eq. (7), once again recovering the expected linear response behaviour close to equilibrium. Moreover, the coefficient describing the linear dependence of the efflux on Δ*µ*_*p*_ near equilibrium is equal to that describing its dependence on Δ*µ*_*D*_, as a direct consequence of the Onsager reciprocity relations along with the fact that the proton flux is equal to the efflux in this model [36].

However, the periplasmic proton concentration has two other critical effects on the efflux rate. These are the Michaelis-Menten-like dependence with affinity *K*_*β*_, and the aforementioned role of [*p*]_*per*_ in setting the MM constant for drug binding. These are both nontrivial effects that depend on [*p*]_*per*_ independently of its role in the electrochemical potential that drives the system away from equilibrium. These three distinct effects give rise to a complicated [*p*]_*per*_-dependence of the efflux. In general, they can oppose one another, making it and difficult to predict, *a priori*, and highly dependent on the values of other parameters, how the efflux will change in response to a change in [*p*]_*per*_.

Finally, just like the proton concentrations, the electric potential difference Δ*V* enters into the equation for *J* not only through the proton motive force. *K*_*G*_ appears on its own in the definition of *K*_*M*_, as well as Eqs. (4) and (5) for the rates of the final, multi-step transition. This dependence is also present only away from equilibrium, vanishing in the linear response regime, where *K*_*G*_ may be taken to 1 as it appears in these contexts.

### C. Tuning the periplasmic pH shifts the optimal *K*_*D*_ range for efficient pumping

A key insight offered by this model of efflux pump kinetics is revealed when examining how the efflux rate varies as a function of the drug dissociation constant, *K*_*D*_, for different values of the periplasmic pH. While *K*_*D*_ influences the value to which the efflux saturates with increasing [*D*]_*in*_, as has been demonstrated in Fig. 3A, each curve in Fig. 4A shows how the efflux varies over a broad range of *K*_*D*_ values. We see that, for a given set of parameter values, the efflux peaks at a particular value of *K*_*D*_ and dies off for both very weakly and very strongly binding molecules. The position of this peak is determined by competing effects of the proton motive force and drug binding affinity.

**FIG. 4.**
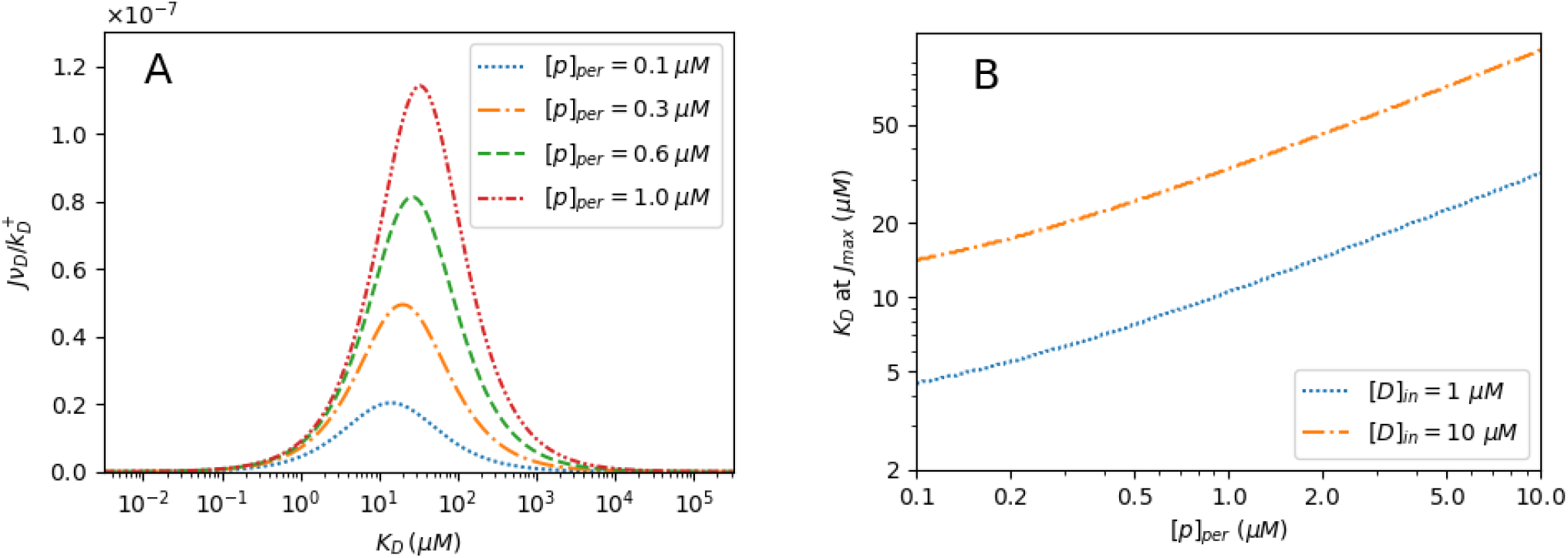
**A**. Dimensionless efflux rate, 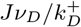, as a function of the drug dissociation constant *K*_*D*_ for varying values of [*p*]_*per*_. The efflux is peaked in a particular *K*_*D*_ range and suppressed on either side of the peak. [*D*]_*in*_ = 10 *µM*. **B**. The value of the drug dissociation constant, *K*_*D*_, at which the efflux rate is maximized, through a range of periplasmic proton concentrations. Other parameter values in both panels are the same as in Fig. 3.

Physically, drugs that bind too weakly (high *K*_*D*_), are likely to unbind from the cavity before the subsequent proton binding can take place. Thus, they cannot proceed to the final transition that sees the drug transported across the membrane. For too strong binding, however, rate 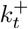 for the final transition that involves the unbinding of the drug into the exit duct is greatly suppressed due to its dependence on *K*_*D*_, as in Eq. (4), limiting the overall rate at which the drug efflux can proceed.

We have established that the efflux rate exhibits MM-like behaviour with respect to the concentration of protons in the cytoplasm. This accounts for the different peak heights between the flux curves in Fig. 4A for different periplasmic proton concentrations. However, these curves also reveal secondary effects of varying [*p*]_*per*_. Namely, through its influence on the Michaelis-Menten constant, *K*_*M*_, changes to [*p*]_*per*_ lead to changes in the values of *K*_*D*_ at which the efflux is maximized. Specifically, in the regime studied, the optimal value of *K*_*D*_ for efflux shifts higher for higher periplasmic proton concentration (lower pH). This shifting effect is visible in Fig. 4A, though somewhat obscured by the logarithmic scale of the horizontal axis. It is emphasized in Fig. 4B, where the *K*_*D*_ value maximizing *J* is shown to shift through a factor of 5-6 as the periplasmic pH varies from 7 to 5. This is understood as a consequence of the fact that increases to [*p*]_*per*_ speed up the 2 → 3 transition towards state 3. The time spent in state 2 is reduced, diminishing the concern that a weak binding drug will detach prematurely, and amplifying the benefits of its ability to readily detach and exit the cell *after* the conformational transition.

The predictions of this model indicate that bacterial populations could, in principle, adapt to drugs with a particular binding affinity to the pump by shifting the periplasmic pH to the value that increases their efflux rate. This suggests a possible mechanism by which multidrug resistance could arise.

In addition to the shifting of the peak, increases to the periplasmic proton concentration grow the overall size of the *K*_*D*_ range at which the efflux is significant. In particular, as seen in Fig. 4A, it extends this range to higher values of *K*_*D*_, representing weaker binding molecules. The high *K*_*D*_ range is precisely where self molecules would be expected to be found, as the cell would be unlikely to have a pump that strongly binds the self molecules it wishes to retain. Thus, increasing [*p*]_*per*_ may increase the risk of extruding self molecules, to detrimental effect, but may still be favourable overall in the presence of a relatively weakly binding drug. Notably, even at high [*p*]_*per*_, the peak is still considerably narrower than that of a proton-independent passive transporter operating under comparable conditions (Supporting Information). That is, in addition to driving the pump and tuning the range of *K*_*D*_ values at which it is most effective, the presence of a proton gradient-dependent transition helps the pump to achieve greater specificity overall.

### D. Specificity tuning behaviour extends to a more detailed model

The three-state model introduced in Sec. IIII A captures the minimal energetic and kinetic features of the pump and allows the derivation of the analytic expression for the efflux, Eq. (7). This serves as a basis for the analysis of the factors affecting efflux pumps’ tunability and identification of the principles of their specificity. However, by grouping multiple steps that include conformational changes, drug unbinding, and proton unbinding into one final transition, it effectively makes the assumption that these molecular processes are all very strongly coupled and occur at a rate determined by the slowest process among them. This model also does not capture the possibility for the drug and proton binding affinities to differ with the conformation of the transporter.

In this section, we study a more complete kinetic scheme which consists of five states, treating the conformational change exposing the drug to the exit duct, drug unbinding to the exit duct, and proton unbinding to the cytoplasm as three separate transitions, each with a distinct forward and reverse rate. This model is depicted in Fig. 5A, where states 1-3 are equivalent to the states of the three-state model. In state 4, the conformational change of the pump has occurred such that the proton binding site now faces the cytoplasm and the interior of the active transporter now opens towards the exit duct, but both the drug and the proton remain bound. In state 5, the drug molecule has been released while the proton remains in place. In the final transition from state 5 back to state 1, the proton is released to the cytoplasm, and the pump, in the absence of a proton responding to the membrane electric potential, promptly returns to its initial conformation. Figure 5B depicts each transition as a directed edge, labelled by its rate.

**FIG. 5.**
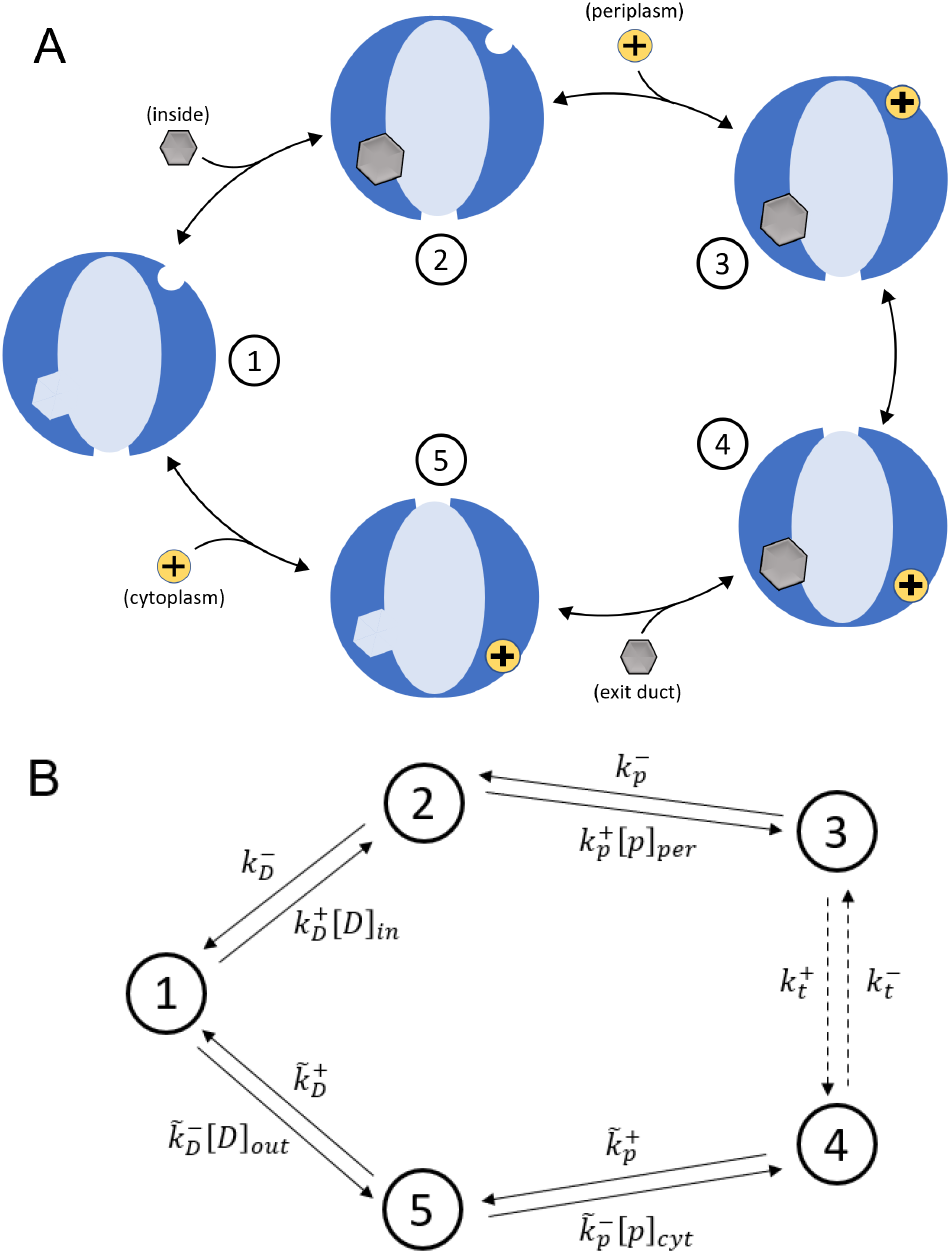
**A**. Five-state model for efflux pump operation. In contrast to the three-state model, the first conformational change, drug unbinding to the exit duct, proton unbinding to the cytoplasm (with return to the initial conformation) are split into separate steps. **B**. The five-state model represented as a directed graph with edges corresponding to transitions. Each transition is labelled with its respective rate, as they arise in the master equation (Supporting Information).

This model is described using the master equation formalism, analogous to Eqs. (1), with a set of differential equations for the probabilities for the system to be found in each of the five states (Supporting Information). The forward and reverse rate constants, 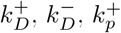 and 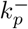 for the transitions between states 1, 2, and 3, are unchanged from the three-state model, along with the dissociation constants *K*_*D*_ and *K*_*p*_ for drug binding from the cytoplasm, and protons from the periplasm, respectively.

The additional flexibility offered by the five-state model, however, allows one to consider precisely how the dissociation constant (and thus, binding energy) for drug molecules may differ in different conformational states of the transporter. That is, the dissociation constant, 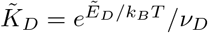, for drug molecules when the pump has undergone a conformational transition and the binding site is exposed to the exit duct is, in general, not equal to *K*_*D*_. The same is true for protons and is modelled by the introduction of 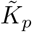 and 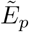, the dissociation constant and binding energy for protons from the cytoplasm. As another additional parameter, we may consider the energy change, *E*_*c*_ ≡ *E*_4_ − *E*_3_, that occurs during the first conformational transition alone. The model is constrained only by the requirement that the system returns to its initial state upon completion of the cycle, so,

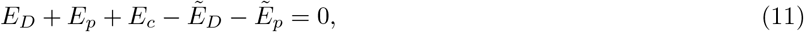

Assuming that the characteristic rates, *r*_*D*_ and *r*_*p*_, and interaction volumes, *ν*_*D*_ and *ν*_*p*_, are the same as above, the forward rate for the transition from state 4 to 5 is 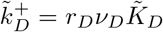 with the reverse rate 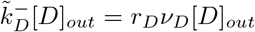.

Similarly, the forward rate for the transition from state 5 to state 1 is 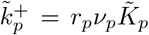, with the reverse rate 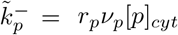. Equation (11) then dictates that *E*_*c*_ is given by

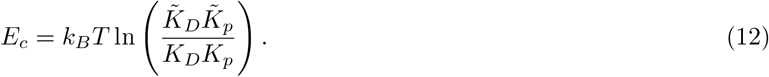

Local detailed balance relations determine the forward and reverse rate constants for the conformational transition between states 3 and 4,

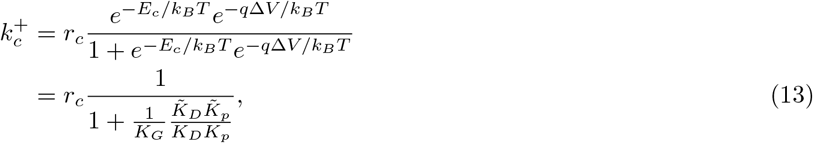

and

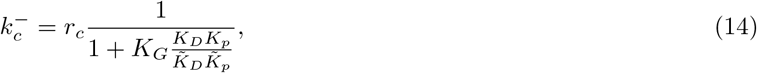

where we introduce *r*_*c*_, a characteristic rate for the conformational change between states 3 and 4. This is a more physically meaningful quantity than its counterpart, *r*_*t*_, in the three-state model, suggesting that moving to five states leads to a more realistic model with less need for fine-tuning. The factor of *K*_*G*_ in 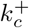 reflects that the electric potential acts on the proton during this step of the cycle, forcing it from an area of high potential (as its binding site is exposed to the periplasm) to an area of lower potential (the cytoplasm), but with it remaining bound throughout this step.

The five-state model does not simplify to provide analytic insights. Therefore, we obtain the efflux rate by numerically solving the system of equations at steady state. Efflux is plotted against *K*_*D*_ in Fig. 6A, showing the same qualitative behaviours seen with the three-state model: maximal efflux at a particular *K*_*D*_ range which shifts higher with increases to [*p*]_*per*_. These results are shown as a contour plot in Fig. 6B, over the periplasmic pH range from 7 to 5.

**FIG. 6.**
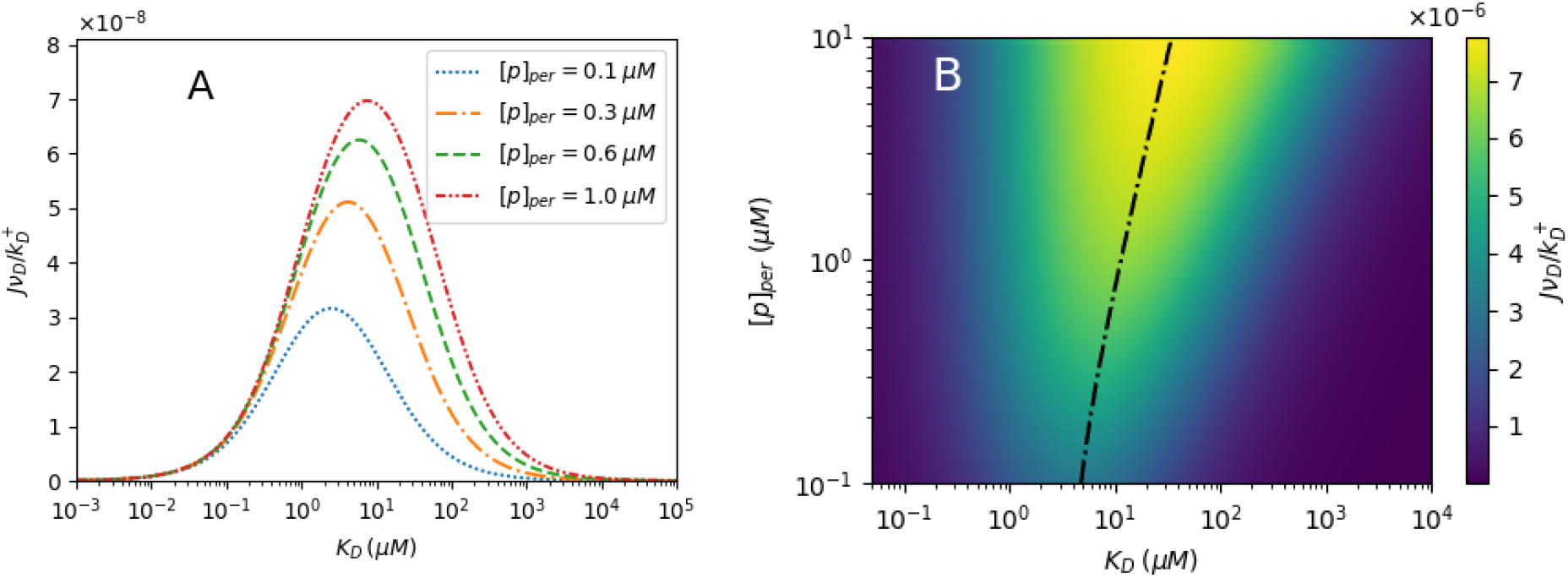
**A**. Dimensionless efflux rate given by the five-state model as a function of the drug dissociation constant, *K*_*D*_, of drug binding from the cytoplasm at different values of the periplasmic proton concentration, [*p*]_*per*_. Similar to the three-state model, we observe maximum efflux within a particular *K*_*D*_ range which shifts towards higher values of *K*_*D*_ with increasing [*p*]_*per*_. **B**. Contour plot showing the variation of the dimensionless efflux rate through the range of periplasmic pH values from 7 to 5. The dash-dotted line represents the value of *K*_*D*_ at which the efflux rate is maximized for each value of [*p*]_*per*_. In both panels, we assume that drug molecules bind less strongly from the outside of the cell: 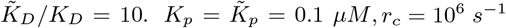, and parameter values are otherwise the same as in Fig. 4.

This model exhibits the peak-shifting effect with varying [*p*]_*per*_ to a similar degree to the three-state model. As seen in Fig. 6, the *K*_*D*_ value for maximum efflux shifts through a factor of 5-6 as the periplasmic pH drops from 7 to 5. For the figure, it is assumed that 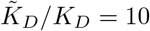, i.e., drugs bind to the transporter with greater affinity when it is in its initial conformation, with the drug binding site facing the interior, than when it is in its second conformation with the binding site facing the exit duct. This would be a feature of a pump that is particularly effective at extruding unwanted molecules from the cell, though the peak-shifting effect is still observed with this assumption relaxed.

We note that we are considering both the tunability of the pump using the five-state model only in a particular region of a very large parameter space, determined based on experimental relevance, but without optimization. If the parameter values were to be optimized to maximize the tunability effect and/or the specificity of the pump, the increased parameter space may allow even better performance to be achieved with the five-state model than with the three-state model.

## III. DISCUSSION

Bacterial efflux pumps are known to play a prominent role in the development of resistance to a broad range of antibiotics. We have studied kinetic models that capture important features of efflux pump operation to elucidate unexpected nonequilibrium effects at play in governing their behaviour. The resulting insights suggest a potential mechanism by which multidrug resistance can arise.

Past studies have utilized kinetic models to help in understanding the translocation process of antibiotics and other molecules within active transporters [10, 26, 27, 50, 53] as well as non-specific transmembrane pores [54]. Most of the published kinetic models are Michaelis-Menten (MM) like in their description of kinetics. Importantly, those describing proton-powered active efflux pumps inherently assume that pumps are driven by the proton motive force and did not consider explicitly the distinct effects of varying the proton concentrations and membrane gradient. Many others considered ATP-powered transporters instead.

We have shown, through analysis using a set of minimal models that explicitly take into account the two separate components of the proton motive force–the membrane potential and the proton concentration gradient–that the operation of cellular efflux pumps exhibits complex and nontrivial dependence on these nonequilibrium electrochemical gradients. It is characterized by behaviour that resembles Michaelis-Menten like kinetics, but with an additional factor resulting from the microscopic reversibility of the process and with effective nonequilibrium Michaelis-Menten constants that differ from expected equilibrium values, which are dictated solely by the binding affinities. These affinities can be modulated by varying the periplasmic proton concentration so as to shift the range of drug dissociation constants (and therefore the drugs themselves) at which the pump operates most effectively. Furthermore, the transporter extrudes molecules over a narrower range of *K*_*D*_ values overall than a passive channel. These effects may offer an explanation as to how bacterial populations can adapt their efflux pumps to particular drugs used in treatment against infections, becoming resistant to antibiotics while maintaining a high degree of specificity.

Until now, common approaches to the inhibition of efflux pumps comprise the targeting of the efflux pumps’ expression, the disruption of the pumps self-assembly process, and the design of new antibiotic molecules that are not recognized by the efflux pumps, as well as new competing substrates to block the pump binding sites [1, 7, 17, 55–58]. In spite of the research efforts, only a limited number of inhibitors are currently in the pre-clinical stage and there are no inhibitor drugs for bacterial efflux pumps in clinical use [55]. The effects of periplasmic pH on pump operation, including the ability to tune their specificity towards molecules in a particular range of binding affinities, may be a consideration as researchers aim to develop treatments for bacterial infections that target the operation of these pumps.

We have characterized the phenomenon of specificity-tuning through alteration of the periplasmic pH via the peak-shifting effect demonstrated in Figs. 4 and 6. An additional effect of increasing the periplasmic proton concentration is reduced specificity, or an overall larger range of *K*_*D*_ values at which the efflux pump throughput is relatively high (Supporting Information). This increases the risk of extruding other cytoplasmic contents, which are expected to be weak binders, characterized by a high *K*_*D*_ value. In the presence of drug molecules, the periplasmic pH can therefore be optimized to best address this trade-off, shifting the efflux peak to target drug molecules, while keeping the pump as specific to the drug as possible.

To emphasize the scope of the proton concentration’s ability to tune efflux behaviour, we consider the linear response regime for the three-state model (Supporting Information). In this case, all thermodynamic forces–the chemical potential differences due to the concentration gradients and the membrane voltage–are small enough relative to the thermal energy that we may Taylor-expand only to linear order in each. To good approximation, the efflux is given by,

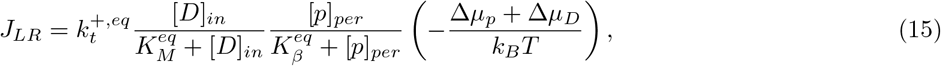

where the superscript *eq* denotes the equilibrium value of each constant, calculated by setting *K*_*G*_ = 1, [*D*]_*out*_ = [*D*]_*in*_, and [*p*]_*cyt*_ = [*p*]_*per*_. Δ*µ*_*p*_ is defined as in Eq. (10) and we define Δ*µ*_*D*_ = *k*_*B*_*T* ln([*D*]_*out*_*/*[*D*]_*in*_). Interestingly, in this regime, while the effect of the proton *gradient* is limited to the linear dependence expressed in Eq. (15), the efflux still depends in an absolute sense on the proton concentration, which scales the overall efflux rate via Michaelis-Menten kinetics, and still plays a role in setting the equilibrium affinity, 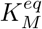.

Another quantity to consider when assessing the efflux pump performance is the entropy production rate. In the absence of external sources, eventually the concentrations of the protons and the drugs will eventually reach a steady state where the probability of the forward cycles is equal to the probability of the backward cycles with 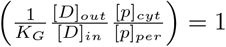 and zero flux. For our models, we have assumed instead that concentrations of drug molecules and protons are kept constant to maintain non-zero efflux. This requires continuous input (and dissipation of energy). This energetic cost can be understood by implicitly assuming that independent translocation processes occur that transport protons and drug molecules back across the membrane to maintain their gradients. Each of these requires a minimum energy input per drug/proton, determined by the change in the chemical potential differences of each in this auxiliary process: *k*_*B*_*T* ln([*D*]_*in*_*/*[*D*]_*out*_) and *k*_*B*_*T* ln([*p*]_*per*_*/*[*p*]_*cyt*_) − *q*Δ*V*, respectively (we note that if the drug concentration is higher outside the cell than inside, ln([*D*]_*in*_*/*[*D*]_*out*_) is negative and the gradient-maintaining process can simply amount to passive transport). Accordingly, the minimal energy dissipation to maintain the steady state non-zero efflux is *J*Δ*F*, where Δ*F* = *k*_*B*_*T* ln([*D*]_*in*_*/*[*D*]_*out*_) + *k*_*B*_*T* ln([*p*]_*per*_*/*[*p*]_*cyt*_) − *q*Δ*V*. Since our models impose a strict one-to-one ratio between the number of drugs extruded and the number of protons transported across the membrane, one can only determine a lower bound on the entropy production rate that is simply proportional to the efflux, *J*. They therefore describe a simple tradeoff relation between pump throughput and dissipation: the greater the efflux, the higher the minimum rate of dissipation necessary to maintain the gradient.

In order to go beyond this simple case, one can augment the kinetic model with the addition of a “futile” cycle, wherein a proton is transported into the cytoplasm, but no drug molecule is extruded (Supporting Information). In this case, the lower bound on the entropy production rate exceeds that predicted from the three- and five-state models, as it must account for the additional proton flux that occurs without any corresponding drug efflux. In incorporating such a futile cycle, however, there is no improvement to the tunability of the pump, and only a minor improvement to the specificity of the efflux pump due to the fact that the futile cycle allows for greater suppression of the efflux at high *K*_*D*_. We do not identify any clear tradeoff relation between specificity and dissipation, distinguishing this study from other works which aim to uncover the constraints thermodynamics place on various measures of precision in cellular processes [59–61].

Conversely, we do find that this model exhibits a greater degree of specificity than a passive channel whose operation is completely independent of the proton electrochemical gradient, suggesting a role is being played by the external nonequilibrium driving force in achieving a certain level of pump performance (Supporting Information).

In this study, we focused on the hitherto unappreciated nonequilibrium effects of the periplasmic pH on the pump throughput. We vary the drug binding affinity and the periplasmic pH as two independent variables, assuming one can be held fixed through a range of values of the other. For practical applications it may be important to consider the direct effects of the pH on the equilibrium binding affinity *K*_*D*_ and *K*_*p*_ [62, 63], and investigate whether this enhances or reduces the specificity and tunability effects outlined in Sec. II.

In summary, we have studied a set of new quantitative dynamical models for bacterial efflux pump active transporters that explicitly take into account the separate components of the proton motive force, membrane potential and the proton gradient. Our model shows that, unlike the commonly held assumptions, the proton concentration gradient does not merely provide the thermodynamic force to drive the pump; it also takes part in determining the pump selectivity. Furthermore, the roles of its two components–the proton concentration gradient and the electric potential gradient across the membrane–enter this determination separately. Thus, modifications to each independently play a role in tuning the efflux mechanism. These results build upon prior understanding of the role that alteration of the free energy landscape characterizing a biological process can play in modulating fluxes [43].

The structure of the model and the main ideas behind its formulation are in line with a number of detailed modeling studies of the Na^+^/glucose cotransporter [24, 40, 41], as well as resistance-nodulation-division pumps such as AcrAB found in *E. Coli* and MexAB in *P. aeruginosa* [14–19]. This new role of the electrochemical force can, in principle, be relevant to other types of ion-driven active transporters as well.

## MATERIALS AND METHODS

### Master equation formalism

The master equations that we use to describe the time evolution of the probability for the system to be in each state, e.g., Eq. (1), can be used to calculate the efflux directly. This probability distribution at any given time is represented by a vector 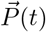 whose dimension is equal to the number of states in the kinetic model. The equations of motion may then be written as a matrix-vector product

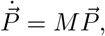

where the each elements *M*_*ij*_ is the rate to transition from state *j* to state *i* and the diagonal elements are set to ensure that Σ _*i*_ *P*_*i*_ = 1 at all times.

Notably, we are interested in the behaviour of the system at steady state, ignoring any changes to the environmental parameters that would affect the matrix *M* on the basis that they would occur on a timescale much longer than the time taken for the probabilities to reach their steady-state values. This amounts to the probability distribution 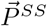 at which 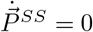, which is equivalent to the normalized eigenvector of *M* associated with the zero eigenvalue.

Equipped with 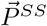 in terms of the model parameters, it is straightforward to calculate the net flux through any given transition. The efflux is identified as the sum of the net fluxes through each transition during which a drug molecule is released to the exit duct. It is, therefore, given by Eq. (6) for the three-state model, as the 2 → 3 transition is the only transition that includes the extrusion of a drug molecule. For a unicyclic model such as those considered here, the net flux through any transition will suffice, since the steady state condition guarantees all these are equal. When considering multicyclic schemes (Supplementary Material), care must be taken in identifying which transitions in the model are associated with the extrusion of drug molecules.

This process for calculating the efflux may be carried out numerically, i.e., by diagonalizing the matrix *M* after the input of numerical values for each parameter. For the three-state model, we were able to do it symbolically as well, leading, after algebraic manipulations, to the derivation of the expression for the efflux rate, Eq. (7), and the associated expressions for the effective affinities.

The identification of the *K*_*D*_ value at which the efflux reaches is maximum, as plotted in Figs. 4B and 6B, was done numerically, by identifying zero-crossings in the numerical derivatives (finite differences) of *J* as a function of *K*_*D*_ for a range of [*p*]_*per*_ values.

## ACKNOWLEDGEMENTS

The work of MG is supported by an NSERC Canada Graduate Scholarship–Doctoral. The work of BSA at Los Alamos National Laboratory, which is operated by Triad National Security, LLC, is supported by National Nuclear Security Administration of U.S. Department of Energy, Contract No. 89233218CNA000001. DS acknowledges support from NSERC and the Canada Research Chairs prgoram. AZ acknowledges the support by NSERC through Discovery Grant.

## References

[1] J. Olivares, A. Bernardini, G. Garcia-Leon, F. Corona, M. B. Sanchez, and J. L. Martinez, The intrinsic resistome of bacterial pathogens, Frontiers in Microbiology 4, 1 (2013).

[2] H. Nikaido, Multidrug resistance in bacteria, Annual Review of Biochemistry 78, 119 (2009).

[3] M. H. Kollef and V. J. Fraser, Antibiotic Resistance in the Intensive Care Unit, Annals of Internal Medicine 134, 298 (2001).

[4] C. J. L. Murray, K. S. Ikuta, F. Sharara, et al., Global burden of bacterial antimicrobial resistance in 2019: a systematic analysis, The Lancet 399, 629 (2022).

[5] N. Goldenfeld and C. Woese, Biology’s next revolution, Nature 445, 2007 (2007).

[6] K. Vetsigian, C. Woese, and N. Goldenfeld, Collective evolution and the genetic code, Proceedings of the National Academy of Sciences of the United States of America 103, 10696 (2006).

[7] P. Hinchliffe, M. F. Symmons, C. Hughes, and V. Koronakis, Structure and operation of bacterial tripartite pumps, Annual Review of Microbiology 67, 221 (2013).

[8] D. Du, Z. Wang, N. R. James, J. E. Voss, E. Klimont, T. Ohene-Agyei, H. Venter, W. Chiu, and B. F. Luisi, Structure of the AcrAB-TolC multidrug efflux pump, Nature 509, 512 (2014).

[9] H. Nikaido, Molecular Basis of Bacterial Outer Membrane Permeability Revisited, Microbiology and Molecular Biology Reviews 67, 593 (2003).

[10] D. G. Thanassi, G. S. Suh, and H. Nikaido, Role of outer membrane barrier in efflux-mediated tetracycline resistance of Escherichia coli, Journal of Bacteriology 177, 998 (1995).

[11] D. Du, X. Wang-Kan, A. Neuberger, H. W. van Veen, K. M. Pos, L. J. Piddock, and B. F. Luisi, Multidrug efflux pumps: structure, function and regulation, Nat. Rev. Microbiol. 16, 523 (2018).

[12] Y. Matsunaga, T. Yamane, T. Terada, K. Moritsugu, H. Fujisaki, S. Murakami, M. Ikeguchi, and A. Kidera, Energetics and conformational pathways of functional rotation in the multidrug transporter acrb, eLife 7, e31715 (2018).

[13] E. M. Darby, E. Trampari, P. Siasat, M. S. Gaya, I. Alav, M. A. Webber, and J. M. A. Blair, Molecular mechanisms of antibiotic resistance revisited, Nat. Rev. Microbiol. 21, 280 (2023).

[14] C. A. Elkins and H. Nikaido, Substrate specificity of the RND-type multidrug efflux pumps AcrB and AcrD of Escherichia coli is determined predominately by two large periplasmic loops, Journal of Bacteriology 184, 6490 (2002).

[15] H. Nikaido, Structure and mechanism of rnd-type multidrug efflux pumps, in Advances in Enzymology and Related Areas of Molecular Biology (John Wiley & Sons, Ltd, 2011) pp. 1–60, https://onlinelibrary.wiley.com/doi/pdf/10.1002/9780470920541.ch1.

[16] A. Yamaguchi, R. Nakashima, and K. Sakurai, Structural basis of RND-type multidrug exporters, Frontiers in Microbiology 6, 1 (2015).

[17] H. Venter, R. Mowla, T. Ohene-Agyei, and S. Ma, RND-type drug efflux pumps from Gram-negative bacteria: Molecular mechanism and inhibition, Frontiers in Microbiology 6, 1 (2015).

[18] S. Minagawa, H. Inami, T. Kato, S. Sawada, T. Yasuki, S. Miyairi, M. Horikawa, J. Okuda, and N. Gotoh, RND type efflux pump system MexAB-OprM of pseudomonas aeruginosa selects bacterial languages, 3-oxo-acyl-homoserine lactones, for cell-to-cell communication, BMC Microbiology 12, 1 (2012).

[19] T. Eicher, M. A. Seeger, C. Anselmi, W. Zhou, L. Brandstätter, F. Verrey, K. Diederichs, J.D. Faraldo-Gómez, and K. M. Pos, Coupling of remote alternating-access transport mechanisms for protons and substrates in the multidrug efflux pump AcrB, eLife 3, 1 (2014).

[20] S. Radestock and L. R. Forrest, The alternating-access mechanism of mfs transporters arises from inverted-topology repeats, Journal of Molecular Biology 407, 698 (2011).

[21] V. Burtscher, K. Schicker, M. Freissmuth, and W. Sandtner, Kinetic models of secondary active transporters, Int. K. Mol. Sci. 20, 5365 (2019).

[22] M. Coincon, P. Uzdavinys, E. Nji, D. L. Dotson, I. Winkelmann, S. Abdul-Hussein, A. Cameron, O. Beckstein, and D. Drew, Crystal structures reveal the molecular basis of ion translocation in sodium/proton antiporters, Nat. Struct. Mol. Biol. 23, 248 (2016).

[23] J. Steiner and L. Sazanov, Structure and mechanism of the mrp complex, an ancient cation/proton antiporter, eLife 2020;9, e59407 (2020).

[24] D. D. Loo, A. Hazama, S. Supplisson, E. Turk, and E. M. Wright, Relaxation kinetics of the Na+/glucose cotransporter, Proceedings of the National Academy of Sciences of the United States of America 90, 5767 (1993).

[25] V. Koronakis, A. Sharff, E. Koronakis, B. Luisi, and C. Hughes, Crystal structure of the bacterial membrane protein TolC central to multidrug efflux and protein export, Nature 405, 914 (2002).

[26] J. Castillo, R. Huan, D. Basilio, A. Das, B. Roux, R. Latorre, F. Bezanilla, and M. Holmgren, Mechanism of potassium ion uptake by the na+/k+-atpase, Nat. Commun. 6, 7622 (2015).

[27] R. Anandakrishnan and D. Zuckerman, Biophysical comparison of atp-driven proton pumping mechanisms suggests a kinetic advantage for the rotary process depending on coupling ratio, PLoS ONE 12, e0173500 (2017).

[28] L. Wang, Z. L. Johnson, M. R. Wasserman, J. Levring, J. Chen, and S. Liu, Characterization of the kinetic cycle of an abc transporter by single-molecule and cryo-em analyses, eLife 2020;9, e56451 (2020).

[29] S. Leipelt and R. Lipowsky, Kinesin’s network of chemomechanical motor cycles, Phys. Rev. Lett. 98, 258102 (2007).

[30] V. Beirbaum and R. Lipowsky, Chemomechanical coupling and motor cycles of myosin v, Biophys. J. 100, 1747 (2011).

[31] S. Kubo, T. Shima, and S. Takada, How cytoplasmic dynein couples atp hydrolysis cycle to diverse stepping motions: Kinetic modeling, Biophys. J. 118, 1930 (2020).

[32] A. Murugan, D. A. Huse, and S. Leibler, Discriminatory proofreading regimes in nonequilibrium systems, Phys. Rev. X 4, 021016 (2014).

[33] D. Kirby and A. Zilman, Proofreading does not result in more reliable ligand discrimination in receptor signaling due to its inherent stochasticity, Proc. Natl. Acad. Sci. 120, e2212795120 (2023).

[34] J. A. Morin, F. J. Cao, J. M. Lazaro, J. R. Arias-Gonzalez, J. M. Valpuesta, J. L. Carrascosa, M. Salas, and B. Ibarra, Mechano-chemical kinetics of dna replication: identification of the translocation step of a replicative dna polymerase, Nucleic Acids Research 43, 3643 (2015).

[35] Y.-S. Song, Y.-G. Shu, X. Zhou, Z.-C. Ou-Yang, and M. Li, Proofreading of dna polymerase: a new kinetic model with higher-order terminal effects, Journal of Physics: Condensed Matter 29, 025101 (2016).

[36] S. R. de Groot and P. Mazur, Non-equilibrium Thermodynamics (Amsterdam North-Holland Pub. Co., 1962).

[37] S. Murakami, R. Nakashima, E. Yamashita, T. Matsumoto, and A. Yamaguchi, Crystal structures of a multidrug transporter reveal a functionally rotating mechanism, Nature 443, 173 (2006).

[38] R. Nakashuima, K. Sakurai, S. Yamasaki, K. Nishino, and A. Yamguchi, Structures of the multidrug exporter acrb reveal a proximal multisite drug-binding pocket, Nature 480, 565 (2011).

[39] E. W. Yu, G. McDermott, H. I. Zgurskaya, H. Nikaido, and D. E. Koshland, Structural Basis of Multiple Drug-Binding Capacity of the AcrB Multidrug Efflux Pump, Science 300, 976 (2003), publisher: American Association for the Advancement of Science.

[40] E. M. Wright, D. D. Loo, and B. A. Hirayama, Biology of human sodium glucose transporters, Physiological Reviews 91, 733 (2011).

[41] L. Parent, S. Supplisson, D. D. Loo, and E. M. Wright, Electrogenic properties of the cloned Na+/glucose cotransporter: I. Voltage-clamp studies, The Journal of Membrane Biology 125, 49 (1992).

[42] Y. C. Kim, M. Wikstrom, and G. Hummer, Kinetic gating of the proton pump in cytochrome c oxidase, Proceedings of the National Academy of Sciences 106, 13707–13712 (2009).

[43] A. I. Brown and D. A. Sivak, Allocating dissipation across a molecular machine cycle to maximize flux, Proceedings of the National Academy of Sciences 114, 11057–11062 (2017).

[44] R. D. Astumian, C. Pezzato, Y. Feng, Y. Qui, P. R. McGonigal, C. Cheng, and J. F. Stoddart, Non-equilibrium kinetics and trajectory thermodynamics of synthetic molecular pumps (2020).

[45] A. Berlaga and A. Kolomeisky, Molecular mechanisms of active transport in antiporters: Kinetic constraints and efficiency, J. Phys. Chem. Lett. 12, 9588 (2021).

[46] H. Qian and H. Ge, Stochastic chemical reaction systems in biology (Springer Nature Switzerland AG, 2021) Chap. Kinetic Rate Equations and the Law of Mass Action.

[47] C. Van den Broeck, Stochastic thermodynamics: A brief introduction, Proceedings of the International School of Physics “Enrico Fermi” 184, 155 (2012).

[48] C. Pezzato, C. Cheng, J. F. Soddart, and R. D. Astumian, Mastering the non-equilibrium assembly and operation of molecular machines, Chem. Soc. Rev. 46, 5491 (2017).

[49] C. C. Su, H. Nikaido, and E. W. Yu, Ligand-transporter interaction in the AcrB multidrug efflux pump determined by fluorescence polarization assay, FEBS Letters 581, 4972 (2007).

[50] K. Nagano and H. Nikaido, Kinetic behavior of the major multidrug efflux pump AcrB of Escherichia coli, Proceedings of the National Academy of Sciences of the United States of America 106, 5854 (2009).

[51] P. Fraņcois, G. Voisinne, E. D. Siggia, G. Altan-Bonnet, and M. Vergassola, Phenotypic model for early t-cell activation displaying sensitivity, specificity, and antagonism, Biophys. J. 110, E888 (2013).

[52] S. Fathi, C. R. Nayak, J. J. Feld, and A. G. Zilman, Absolute ligand discrimination by dimeric signaling receptors, Biophysical Journal 111, 917 (2016).

[53] D. Fange, K. Nilsson, T. Tenson, and M. Ehrenberg, Drug efflux pump deficiency and drug target resistance masking in growing bacteria, Proceedings of the National Academy of Sciences of the United States of America 106, 8215 (2009).

[54] M. Ceccarelli, A. V. Vargiu, and P. Ruggerone, A kinetic Monte Carlo approach to investigate antibiotic translocation through bacterial porins, Journal of Physics Condensed Matter 24, 10.1088/0953-8984/24/10/104012 (2012).

[55] T. J. Opperman and S. T. Nguyen, Recent advances toward a molecular mechanism of efflux pump inhibition, Frontiers in Microbiology 6, 1 (2015).

[56] T. Rushikesh, N. Mahey, N. Chandal, D. K. Verma, M. Jangra, et al., A microbe-derived efflux pump inhibitor of the resistance-nodulation-cell division protein restores antibiotic susceptibility in escherichia coli and pseudomonas aeruginosa, ACS Infectious Diseases 8, 255 (2022).

[57] A. P. MacGowan, M. Attwood, A. R. Noel, R. Barber, Z. Aron, et al., Exposure of escherichia coli to antibiotic-efflux pump inhibitor combinations in a pharmacokinetic model: impact on bacterial clearance and drug resistance, Journal of Antimicrobial Chemotherapy (2023).

[58] T.-V. Phan, V.-T.-V. Nguyen, C.-H.-H. Nguyen, T.-T. Vu, T.-D. Tran, et al., Discovery of acrab-tolc pump inhibitors: Virtual screening and molecular dynamics simulation approach, Journal of Biomolecular Structure and Dynamics (2023).

[59] A. H. Lang, C. K. Fisher, T. Mora, and P. Mehta, Thermodynamics of statistical inference by cells, Phys. Rev. Lett. 113, 148103 (2014).

[60] J. D. Mallory, A. B. Kolomeisky, and O. A. Igoshin, Trade-offs between error, speed, noise, and energy dissipation in biological processes with proofreading, J. Phys. Chem. 123, 4718 (2019).

[61] Q. Yu, J. D. Mallory, A. B. Kolomeisky, J. Ling, and O. A. Igoshin, Trade-offs between speed, accuracy, and dissipation in trnaile aminoacylation, J. Phys. Chem. Lett. 11, 4001 (2020).

[62] L. J. Olson, O. Hindsgaul, N. M. Dahms, and J.-J. P. Kim, Structural insights into the mechanism of ph-dependent ligand binding and release by the cation-dependent mannose 6-phosphate receptor*, Journal of Biological Chemistry 283, 10124 (2008).

[63] I. Jarmoskaite, I. AlSadhan, P. P. Vaidyanathan, and D. Herschlag, How to measure and evaluate binding affinities, eLife 9, e57264 (2020).

