## Supporting Information for "Specificity and tunability of efflux pumps: a new role for the proton gradient?"

### Supporting Information for
#### Specificity and tunability of efflux pumps: a new role for the proton motive force?

Matthew Gerry, Duncan Kirby, Boian S. Alexandrov, Dvira Segal, and Anton Zilman

##### A. Michaelis-Menten constant for the efflux rate

As discussed in Sec. 1B of the main text, the efflux rate exhibits Michaelis-Menten (MM)-like behaviour with respect to the concentrations of both drug molecules inside the cell and protons in the cytoplasm, with effective MM constants $K_M$ and $K_\beta$, respectively. This is described precisely in Eqs. (7)-(9) of the main text, where expressions for the efflux itself and $K_\beta$ are given, along with an approximate expression for $K_M$ that holds well in the experimentally relevant regime. For completeness, we state here the exact expression for the MM-constant, $K_M$ associated with the drug concentration:

$$K_M = K_D \frac{\frac{r_t}{r_D}\nu_p[p]_{per}K_G + C + \frac{[D]_{out}[p]_{cyt}}{K_D K_p}\left(\frac{r_t}{r_p}\nu_D K_D + \frac{r_t}{r_D}\nu_p K_p\left(1 + \frac{[p]_{per}}{K_p}\right)\right)}{\frac{[p]_{per}}{K_p}\left(1 + \nu_D\nu_p K_D K_p K_G\right) + C}, \tag{1}$$

where $C = 1 + \nu_D K_D K_G(\frac{r_t}{r_p} + \nu_p K_p)$.

##### B. Efflux pumps in the linear response regime

Close to thermal equilibrium, a nonequilibrium flux such as the efflux of our pumps should have linear dependence on the thermodynamic affinities that drive it: in this case, the proton motive force $\Delta\mu_p = k_B T \ln([p]_{cyt}/[p]_{per}) + q\Delta V$ and the chemical potential difference of drug molecules $\Delta\mu_D = k_B T \ln([D]_{out}/[D]_{in})$. Referring to Eq. (7) in the main text describing the mean efflux for the three-state model, this dependence arises in the factor inside the brackets, which, expanded to linear order in the thermodynamic forces, is given by

$$\left(1 - \frac{1}{K_G}\frac{[D]_{out}}{[D]_{in}}\frac{[p]_{cyt}}{[p]_{per}}\right) = \left(1 - e^{\frac{q\Delta V}{k_B T}}\frac{[D]_{out}}{[D]_{in}}\frac{[p]_{cyt}}{[p]_{per}}\right) \approx -\frac{\Delta\mu_p + \Delta\mu_D}{k_B T}. \tag{2}$$

Here, the membrane voltage and proton chemical potential gradient each factor in only as a component of the proton motive force, contrasting the more complex behaviour highlighted in the general case. That is, the efflux depends linearly on the chemical potential gradient and the voltage with the same coefficient. It is given by

$$J_{LR} = k_t^{+,eq}\frac{[D]_{in}}{K_M^{eq} + [D]_{in}}\frac{[p]_{per}}{K_\beta^{eq} + [p]_{per}}\left(-\frac{\Delta\mu_p + \Delta\mu_D}{k_B T}\right), \tag{3}$$

where the prefactor includes equilibrium values of the constants $K_M$, $K_\beta$, and $k_t^+$, each denoted with the superscript *eq*. These are derived by setting $K_G = 1$ and assuming each concentration varies negligibly between the separate regions, i.e., $[D]_{out} \approx [D]_{in}$ and $[p]_{cyt} \approx [p]_{per}$, eliminating any dependence of the efflux on thermodynamic forces beyond linear order. We note, however, that the linear response coefficient does depend on the overall values of concentrations of both drug molecules and protons. This is demonstrated in Fig. S1, where the dimensionless efflux is shown as a function of the proton motive force for three different values of $[D] \equiv [D]_{in} \approx [D]_{out}$. Even at linear response, the nontrivial role played by the proton concentration in tuning the efflux behaviour is present.

Moreover, the efflux exhibits the exact same dependence on the drug chemical potential gradient as it is does on the proton electrochemical gradient, as a direct reflection of the expected Onsager reciprocal relations along with the fact that this model only has a single flux, i.e., proton influx and drug efflux are equivalent.

##### C. Determination of parameter values

While our main focus is on the relationships between the various parameters describing the model, we determine experimentally relevant numerical values and ranges for these parameters for the purpose of plotting our results. Experiments on efflux pumps have shown that $K_D$ falls in the range of $5 - 74\ \mu M$ [1]. The interaction volume is then taken to $1/\nu_D = 1\ M$, assuming an interaction length scale of about 10 $nm$. $r_D$ is set to $10^8\ s^{-1}$, setting values of $k_D^+$ and $k_D^-$ consistent with diffusion-limited drug-pump interactions [2].

The determination of the analogous quantities for proton binding is less straightforward, due in part to the variability of proton binding affinity. For example, estimates for $K_p$ of aspartic acid, which makes up the most relevant residues for proton binding to the pump in the case of the AcrAB-TolC transporter [3], range as high as about $10^2\ \mu M$ [4] and as low as $10^{-3}\ \mu M$ [5]. We choose $K_p = 0.1\ \mu M$ as a representative value and note that our qualitative findings are not so sensitive to this choice as long as we remain outside the regime that $r_t\nu_p K_p K_G/r_D \approx 1$ (see Sec. D). The identification of $\nu_p$ is similarly challenging; a value of $\nu_p = 10^{-6}\ M^{-1}$ is consistent with proton binding energy on the order of $-10k_BT$, which we set as a reasonable guideline based on measurements of other proteins [6]. We also set $r_p = 10^{14}\ s^{-1}$ to obtain $k_p^+$ of about $10^8\ s^{-1}M^{-1}$, again, consistent with measurements of other proteins [7].

$K_G$ is set to $10^2$, reflecting a membrane potential on the order of $-100\ mV$ at $T = 298\ K$. Finally, we take $r_t = 10^{17} s^{-1}$, making $k_t^+$ comparable to $k_p^-$, since proton unbinding is expected to be the slowest of the multiple processes that occur in this transition. We stress that $r_t$ does not have a clear physical interpretation, but is rather a free parameter that arises as a result of the simplicity of the model, and acknowledge that its value is chosen with the values of other independent parameters in mind, making it, to a degree, regime-dependent. It is replaced by more physically meaningful quantities when the strong assumptions of cooperativity are relaxed.

##### D. Dependence on the proton binding affinity

As discussed in the main text, the proton binding affinity, $K_p$, is a difficult parameter to estimate. Its value is highly context dependent and may not resemble values determined by experiments on the amino acids making up the binding sites on the transporter. For example, the residues associated with the proton binding sites on the AcrAB-TolC transporter consist of aspartic acid [3]. Estimates of $K_p$ for aspartic acid range as high as about $10^2\ \mu M$ [4] and as low as $10^{-3}\ \mu M$ [5]. To explore the effects of varying $K_p$ on the results discussed in the main text, we plot in Fig. S2 the efflux predicted by the three-state model as a function of both $K_D$ and $[p]_{per}$ for three choices of $K_p$: 1 $nm$, corresponding to the regime in which protons bind strongly to the transporter, 0.1 $\mu M$, representing the intermediate regime chosen for all other figures, and 10 $\mu M$, representing the regime of very weak binding.

Only in this latter, weak binding regime is the peak-shifting effect discussed in the main text severely diminished (though it still arises at sufficiently low periplsamic pH). To understand this we repeat here the approximate expression for $K_M$,

$$K_M \approx K_D \frac{\frac{r_t}{r_D}\nu_p K_G [p]_{per} + C}{\frac{[p]_{per}}{K_p} + C}, \tag{4}$$

and note that $K_p \approx 10\ \mu M$ achieves the condition that $r_t\nu_p K_p K_G/r_D \approx 1$. In this very special case, the effective affinity is equal to the equilibrium dissociation constant $K_D$ to good approximation, regardless of the value of $[p]_{per}$.

Thus, the peak-shifting effect is present as long as the proton dissociation constant is relatively low when compared with the drug dissociation constant $K_D$. In this case, the degree to which the $2 \rightarrow 3$ transition in the kinetic scheme is biased towards state 3 is sensitive to changes to $[p]_{per}$ values within the physiological range. This, in turn, influences whether the transporter more effectively pumps stronger binding drugs (which are less likely to detach before a proton has a chance to bind) or weaker binding drugs (which more readily detach towards the exit duct once a proton has bound and the conformation of the pump has changed).

Furthermore, it is evident in Fig. S2 that the efflux tends to be peaked at higher values of $K_D$ overall when $K_p$ is lower. This is because lowering $K_p$ has an effect similar (though not equivalent) to increasing $[p]_{per}$, biasing the $2 \rightarrow 3$ towards state 3, leading the system to favour weaker binding drugs that detach more readily. $K_p$ can therefore be seen as a parameter that both influences the $K_D$ values at which the pump is most effective, and modulates the ability of $[p]_{per}$ to do the same thing.

Finally, increases to $K_p$ increase the efflux overall in the regime studied, as protons, too, detach from the transporter more readily after the conformational transition.

##### E. Master equations for the five-state model

The five-state model considered in Sec. 1D is governed by a set of linear, homogeneous master equation analogous to those for the three-state model, given in Eq. (1) of the main text. With the rate constants defined in the main text, the master equations are given here for completeness:

$$\begin{aligned}
\dot{P}_1 &= -(k_D^+[D]_{in} + \tilde{k}_p^-[p]_{cyt})P_1 + k_D^- P_2 + \tilde{k}_p^+ P_5\\
\dot{P}_2 &= k_D^+[D]_{in}P_1 - (k_D^- + k_p^+[p]_{per})P_2 + k_p^- P_3\\
\dot{P}_3 &= k_p^+[p]_{per}P_2 - (k_p^- + k_c^+)P_3 + k_c^- P_4\\
\dot{P}_4 &= k_c^+ P_3 - (k_c^- + \tilde{k}_D^+)P_4 + \tilde{k}_D^-[D]_{out}P_5\\
\dot{P}_5 &= \tilde{k}_p^-[p]_{cyt}P_1 + \tilde{k}_D^+ P_4 - (\tilde{k}_D^-[D]_{out} + \tilde{k}_p^+)P_5.
\end{aligned} \tag{5}$$

The efflux is calculated on the basis of a steady state solution of these equations (all $\dot{P}_i = 0$). This is carried out numerically by identifying the eigenvector of the associated $5 \times 5$ matrix corresponding to the zero eigenvalue.

While it may be tempting to assume so, we note that the five-state model is not equivalent to the three-state model of Sec. 1A of the main text in any valid limit. Such an equivalence would be based on the grouping of states 3, 4 and 5 into one so-called "metastate", characterized by the pump being in *one of* states 3, 4, or 5, but without specifying which. However, this is not equivalent to the original three-state kinetic scheme, since the condition of Markovianity is broken here: the rates to transition to state 1, completing the cycle, or back to state 2, depend on the state *from* which the system entered this "metastate". Markovianity is recovered only in the case that all transition rates between states 3 and 4, and 4 and 5, are much faster than the rest of the rates in the problem. However, this cannot be achieved without violating some of the constraints on how these rates compare, set by their definitions. As such, while the three-state and five-state models presented so far are closely related, they should be understood as two distinct and alternate kinetic schemes for pump operation.

##### F. Multicyclic model with a futile cycle

The three- and five-state kinetic schemes discussed in the main text exemplify how efflux pumps respond to changes in the parameters governing their operation, such as drug binding affinity and periplasmic proton concentration. However, upon inspection of the cycles depicted in Figures 2 and 5 of the main text, one may note that the pump invariably operates with perfect "chemical efficiency", $J/J_p = 1$ where $J_p$ is the flux of protons into the cytoplasm. That is, due to the unicyclic nature of the models, for every proton transported to the cytoplasm, exactly one drug molecule is removed from the cell.

We wish to consider the situation in which some protons are able to leak through the inner membrane under the proton motive force without any drug molecules being pumped out. This may serve as an additional element of the explanation as to why the cell does not simply maximize the proton concentration in the periplasm, despite this leading to higher efflux overall. We augment the five-state kinetic scheme by adding an additional two states to obtain a multicyclic network, as observed in Fig. S3A, with the network depiction of the model annotated with transition rates in Fig. S3B. The prior five-state "pump" cycle is present, consisting of the identical states 1-5, however, the system can also transition from state 1 to the new states 6 and 7, amounting to a new "waste" cycle. In this cycle, following Figure S3A along the clockwise direction, a proton first binds to the pump in the absence of any drug molecule. The conformational change subsequently takes place. Then, the proton unbinds, entering the cytoplasm. This constitutes a proton being wasted: it is transported across the inner membrane without the associated extrusion of a drug molecule.

This augmentation to the model amounts to the introduction of yet more parameters. To obtain the master equations for this model, we begin with those for the five-state model, Eq. (5), but relabel $K_p \to K_{p,pump}$, and $\tilde{K}_p \to \tilde{K}_{p,pump}$, and add the subscript "*pump*" to rate constants that depend on these quantities, as introduced in Sec. 1D of the main text. This is because we now allow, in principle, the relevant binding affinities to take on different values in the waste cycle: $K_{p,waste}$ and $\tilde{K}_{p,waste}$. That is, the binding affinity of protons, whether from the periplasm or cytoplasm, may differ depending on whether binding occurs in the presence or absence of a bound drug molecule. We define the transition rate constants for this cycle as $k^+_{p,waste} = k^+_{p,pump}$ (and analogously $\tilde{k}^-_{p,waste} = \tilde{k}^-_{p,pump}$), but the unbinding rates differ:

$$k^-_{p,waste} = r_p \nu_p K_{p,waste} \text{ and } \tilde{k}^+_{p,waste} = r_p \nu \tilde{K}_{p,waste}. \tag{6}$$

Accordingly, the rates for the conformational transition differ as well when taking place during the waste cycle:

$$k^+_{c,waste} = r_t \frac{1}{1 + \frac{1}{K_G}\frac{\tilde{K}_{p,waste}}{K_{p,waste}}}, \tag{7}$$

and,

$$k_{c,waste}^{-} = r_t \frac{1}{1 + K_G \frac{K_{p,waste}}{\tilde{K}_{p,waste}}}. \tag{8}$$

The master equations match Eq. (5) for the populations of the states that occur only as part of the pump cycle (i.e., states 2 through 5). However, the addition of the waste cycle brings some new contributions to $\dot{P}_1$ and new equations for $\dot{P}_6$ and $\dot{P}_7$. We have,

$$\begin{aligned}
\dot{P}_1 &= -(k_D^+[D]_{in} + \tilde{k}_D^-[D]_{out} + k_{p,waste}^+[p]_{per} + \tilde{k}_{p,waste}^-[p]_{cyt})P_1 + k_D^- P_2 + \tilde{k}_D^+ P_5 + k_{p,waste}^- P_6 + \tilde{k}_{p,waste}^+ P_7 \\
\dot{P}_6 &= k_{p,waste}^+[p]_{per} P_1 - (k_{p,waste}^- + k_{c,waste}^+)P_6 + k_{c,waste}^- P_7 \\
\dot{P}_7 &= \tilde{k}_{p,waste}^-[p]_{cyt} P_1 + k_{c,waste}^+ P_6 - (\tilde{k}_{p,waste}^+ + k_{c,waste}^-)P_7.
\end{aligned} \tag{9}$$

In Fig. S4A, we plot the efflux of the seven-state model as a function of $K_D$ for various values of $[p]_{per}$, as in the main text for the three- and five-state models. Inspecting these curves, one can observe the same peak-shifting and broadening effects as the periplasmic pH drops. Additionally, at higher $[p]_{per}$ values, there is an overall suppression of the efflux when compared with the unicyclic models. This can be attributed to the presence of the futile cycle in the model–a high value of $[p]_{per}$ is understood to lead the system to spend more of its time in this cycle, and thus less time actively extruding drug molecules from the cell. Thus, the addition of a futile cycle to the model detracts from the apparent finding that increasing the periplasmic proton concentration always increases the efflux, introducing an additional consideration to the process of optimizing the periplasmic pH.

Correspondingly, one can evaluate the chemical efficiency, $J/J_p$, of the pump when described by a multicyclic model: that is, how many protons are transported into the cell on average *per* drug molecule extruded. As depicted in Fig. S4B, this quantity drops as $[p]_{per}$ rises and the pump shifts towards futile cycling. However, there is also a drop-off in chemical efficiency seen above a certain value of $K_D$, regardless of $[p]_{per}$. This is because at very high $K_D$, drugs often cannot stay bound to the pump long enough for the pumping cycle to proceed. This again leads the futile cycle to dominate the pump's operation. We note that this drop off corresponds to the range in $K_D$ at which the overall efflux drops off. Conversely, in the regime of very low $K_D$, where the efflux is also low, the chemical efficiency is relatively high.

Another consequence of moving to a multicyclic model is that the calculation of the minimum entropy production rate is less straightforward. It is no longer directly proportional to the efflux, as there are now contributions from the futile cycle. Directly from the master equations and steady-state probability distribution, the minumum entropy production rate required to maintain the drug and proton gradients is given by the relation:

$$\dot{S} = k_B \sum_{i \neq j} P_i k_{ji} \ln\left(\frac{k_{ji}}{k_{ij}}\right), \tag{10}$$

where the summation is over pairs of states states in the model, $P_i$ is the steady-state population of site $i$, and $k_{ji}$ is the overall transition rate from site $i$ to site $j$ (e.g., $k_{21} = k_D^+[D]_{in}$, $k_{43} = k_c^+$, etc.). We plot in Fig. S4C and D a dimensionless version of this quantity defined as the entropy production rate normalized by the heat dissipated as one proton moves down the electric potential gradient, as well as $k_D^+/\nu_D$, the timescale associated with drug diffusion to the binding site once it is inside the interaction volume,

$$\dot{\Sigma} = \frac{T\dot{S}}{q\Delta V} \frac{\nu_D}{k_D^+}. \tag{11}$$

While comparable minimum entropy production is observed at $K_D = 10\ \mu M$ for both the five- and seven-state model, for weakly binding drugs ($K_D = 100\ \mu M$), the multicyclic model exhibits a much higher minimum entropy production rate than the five-state unicyclic model. This is particularly true as the proton concentration in the periplasm increases. We can understand this as a consequence of the fact that, for both models, the higher-$K_D$ regime amounts to a suppression of the efflux. However, for the unicyclic models this is inextricably linked to a reduction of the minumum entropy production rate. The same cannot be said when the waste cycle in included, giving the system a pathway to allow protons to be transported down their electrochemical gradient even when the drug efflux is suppressed. Notably, in our calculations, we have set the drug gradient to zero: $[D]_{in} = [D]_{out}$. Thus, all contributions to $\dot{\Sigma}$ arise from proton transport and the two cycles give rise to an equal contribution. If one were to suppose a higher drug concentration outside the cell, the contribution to $\dot{\Sigma}$ from the pumping cycle would be partially offset by the fact that drug molecules are being pumped against their concentration gradient, amplifying the tendency of the multicyclic model to show a higher minumum entropy production rate in the regime characterized by a high degree of waste.

Finally, we wish to comment on the predictions the multicyclic model makes for the specificity on the pump, namely, whether the increased dissipation is associated with any improved specificity. Studies have identified constraints on various measures of precision in cellular processes placed by thermodynamics [8–10]. Here, we do observe that the efflux is restricted to a slightly narrower range with the seven-state model than with the five-state model, once again due to the tendency of the system to spend a considerable amount of time in the waste cycle when drug efflux is suppressed. As a result, the efflux begins to drop off at somewhat lower values of $K_D$. This effect is seen by inspecting Fig. S4A, in comparison to Fig. 6A of the main text, however, in the parameter regime studied, it is relatively minor.

##### G. Comparison to a proton-independent channel

We emphasize the advantages that the proton-dependent cycle we introduce in the main text offers with regards to specificity and tunability by considering an analogous model which does not depend on proton concentrations, and instead acts as a passive transporter of drug molecules. We do this in the simplest possible way, augmenting the three-state model by removing proton concentration dependence from all transition rates and eliminating the proton-binding step. This yields a two state model wherein a drug may bind to the transporter (analogous to the $1 \to 2$ transition in the standard, three-state model), and then a second combined step consists of a conformational change, drug unbinding, and return to the original conformation. The master equations describing this model are

$$\begin{aligned}\dot{P}_1 &= -(k_D^+[D]_{in} + k_t^{*,-}[D]_{out})P_1 + (k_D^- + k_t^{*,+})P_2 \\ \dot{P}_2 &= (k_D^+[D]_{in} + k_t^{*,-}[D]_{out})P_1 - (k_D^- + k_t^{*,+})P_2,\end{aligned} \tag{12}$$

where $k_D^+$ and $k_D^-$ are identical to that for the three-state model, but we introduce new rates,

$$k_t^{*,+} = r_t^* \frac{\nu_D K_D}{1 + \nu_D K_D},\ k_t^{*,-} = r_t^* \frac{\nu_D}{1 + \nu_D K_D}, \tag{13}$$

to reflect the distinct processes that occur during the associated transition. Note that there is no dependence on $K_G$ as the lack of involvement from any protons leaves no component that interacts with the electric field.

The efflux through this passive channel is given by

$$J = k_t^{*,+} \frac{[D]_{in}}{K_M^* + [D]_{in}} \left(1 - \frac{[D]_{out}}{[D]_{in}}\right), \tag{14}$$

taking a highly analogous form to the expression for the three-state model, but with no dependence on proton concentrations and a much simpler effective affinity:

$$K_M^* = K_D \left(1 + \frac{r_t^*}{r_D} \frac{1}{1 + \nu_D K_D} \left(1 + \frac{[D]_{out}}{K_D}\right)\right). \tag{15}$$

Figure S5 shows the efflux for this model, with $[D]_{out} = 0.1\ \mu M$. This value, lower than those used in the main text, allows there to be nonzero efflux with $[D]_{in}$ values comparable to those in the main text, since in this case, a drug concentration gradient is needed as there are no other driving forces. Also, $r_t^*$ is set at $10^8\ s^{-1}$, as needed to give a physically realistic transition rates. Accordingly, for experimentally relevant values of $r_D$, $r_t^*$, and $K_D$, $K_M^*$ is well approximated by $K_D$ (at least as an order-of-magnitude estimation), provided it is not that case that $[D]_{out} \gg K_D$. Thus, outside this special case, the drug binding affinity to the pump essentially sets the effective affinity in determining the efflux rate, representing the loss of nontrivial nonequilibrium effects we have identified in the other models of efflux pumps we have considered.

Inspecting Fig. S5, it is clear that the lack of the proton-dependent processes leads to pump operation exhibiting a significantly lesser degree of specificity. The efflux vs. $K_D$ curve maintains non-negligible values up to much higher $K_D$, highlighting the role of the proton gradient not only in tuning the position of the efflux peak, but also in moderating its width. This can be attributed to the manner in which the three-state pump limits the efflux of weaker-binding drugs: the finite rate at which the proton binding step occurs gives the drug molecule time to unbind back into the interior of the cell before the conformational change takes place. In the model considered here, no such mechanism is in place–the conformational change may occur as soon as the drug binds. This distinction between the two models is an example of improved performance that comes along with an additional source of entropy production (i.e. the transport of protons down their electrochemical gradient).

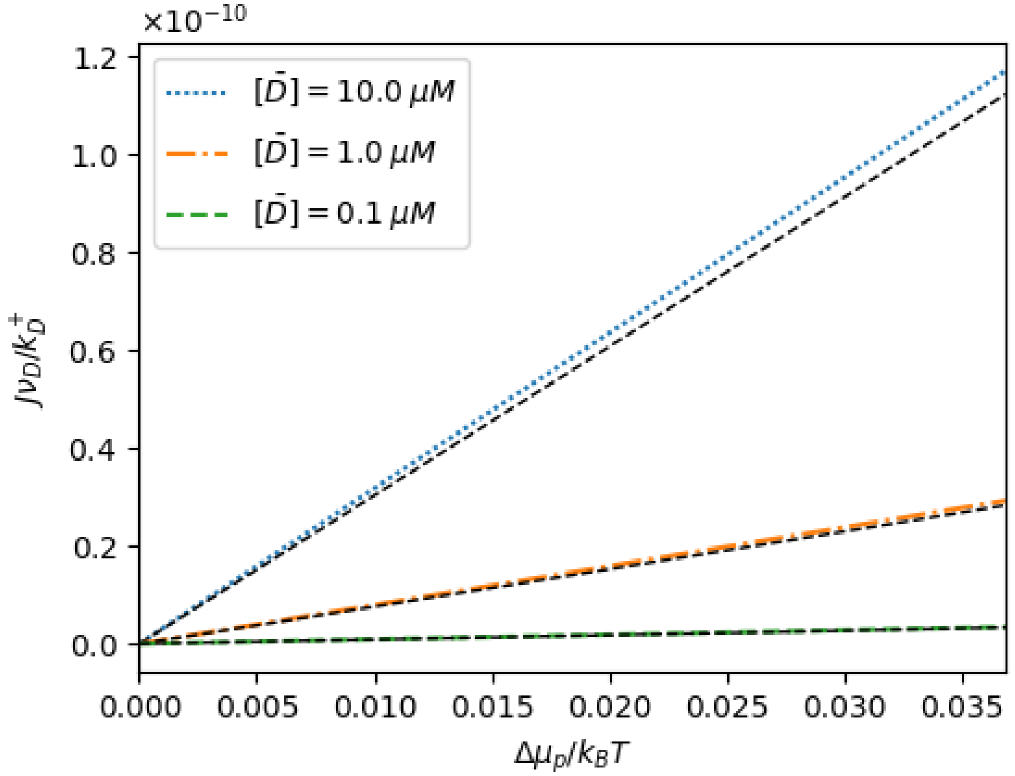


FIG. S1. The dimensionless efflux rate for the three-state model near chemical equilibrium. The colour lines represent the approximation of linear response predictions, with the black dashed lines showing the exact values. For this figure, $\Delta V = 0$, so the quantity plotted on the horizontal axis, $\Delta\mu_p/k_BT$, is equivalent to $\ln([p]_{per}/[p]_{cyt})$, the log-ratio of concentrations on either side of the membrane. This takes on values less than 0.1 representing the linear response regime, with the approximation holding best for the lowest values. As predicted, the slope of the line depends on the drug concentration $[\bar{D}]$, taken to be the same inside and outside the cell ($\Delta\mu_D = 0$). $K_D = 10\ \mu M$, the proton concentration in the cytoplasm is held fixed at $[p]_{cyt} = 0.1\ \mu M$, and the remaining parameter values are the same as in the figures in the main text: $r_D = 10^8\ s^{-1}$, $r_p = 10^{14}\ s^{-1}$, and $r_t = 10^{17}\ s^{-1}$; $\nu_D = 1\ M^{-1}$ and $\nu_p = 10^{-6}\ M^{-1}$; $K_p = 0.1\ \mu M$.

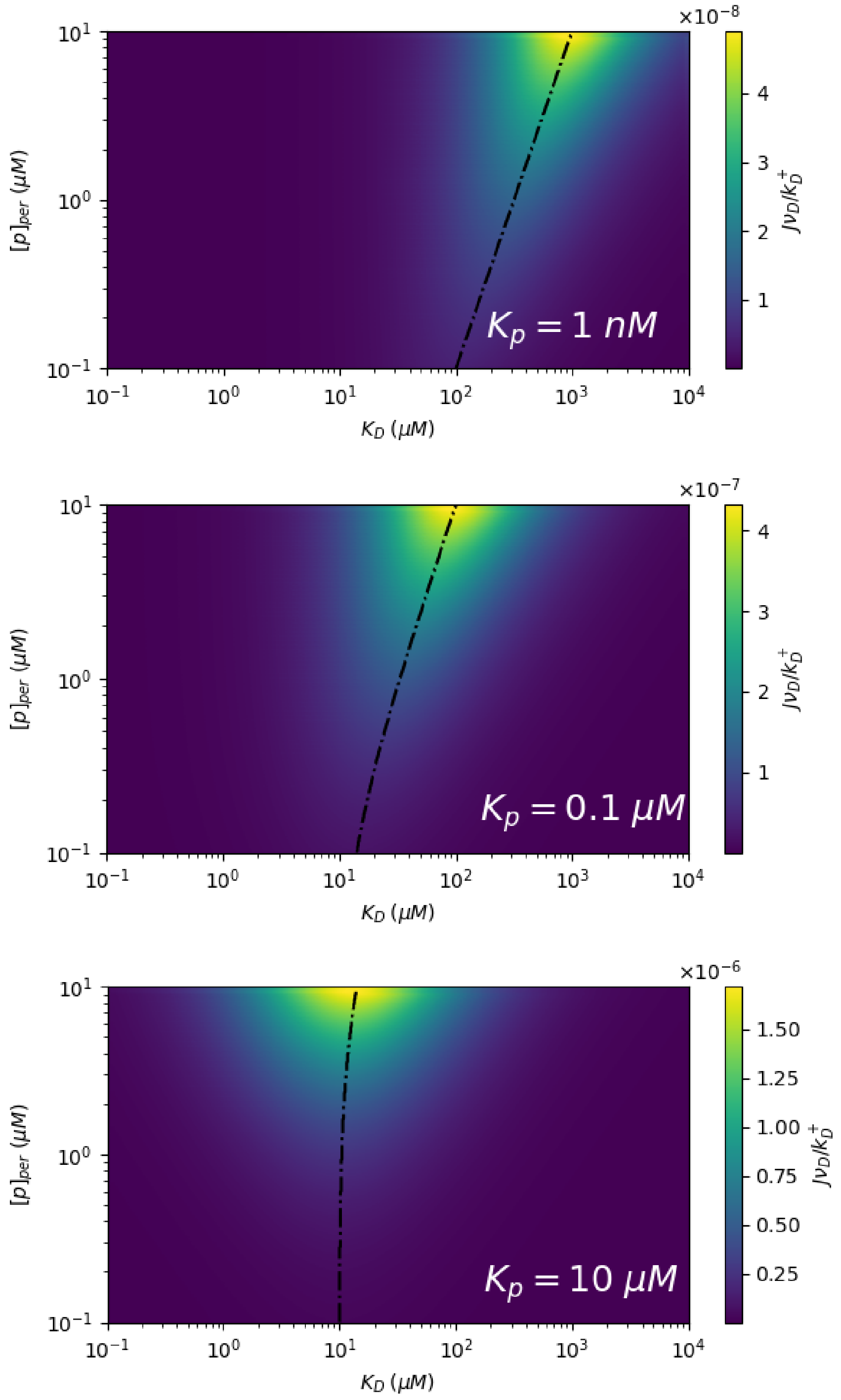


FIG. S2. Contour plots showing the non-dimensional efflux as a function of both $K_D$ and $[p]_{per}$ for the three-state model at three different values of $K_p$, representing strong (upper), intermediate (middle), and weak binding (lower). The dash-dotted curve indicates the point at which the maximum efflux is reached at each value of $[p]_{per}$. The effect of peak-shifting with increasing $[p]_{per}$ is observed dramatically at strong and intermediate proton binding, though diminished at weak binding. Parameter values other than $K_p$ are the same as in the figures of the main text: $r_D = 10^8\ s^{-1}$, $r_p = 10^{14}\ s^{-1}$, and $r_t = 10^{17}\ s^{-1}$; $\nu_D = 1\ M^{-1}$ and $\nu_p = 10^{-6}\ M^{-1}$; $K_G = 100$; $[D]_{in} = [D]_{out} = 10\ \mu M$ and $[p]_{cyt} = 0.1\ \mu M$.

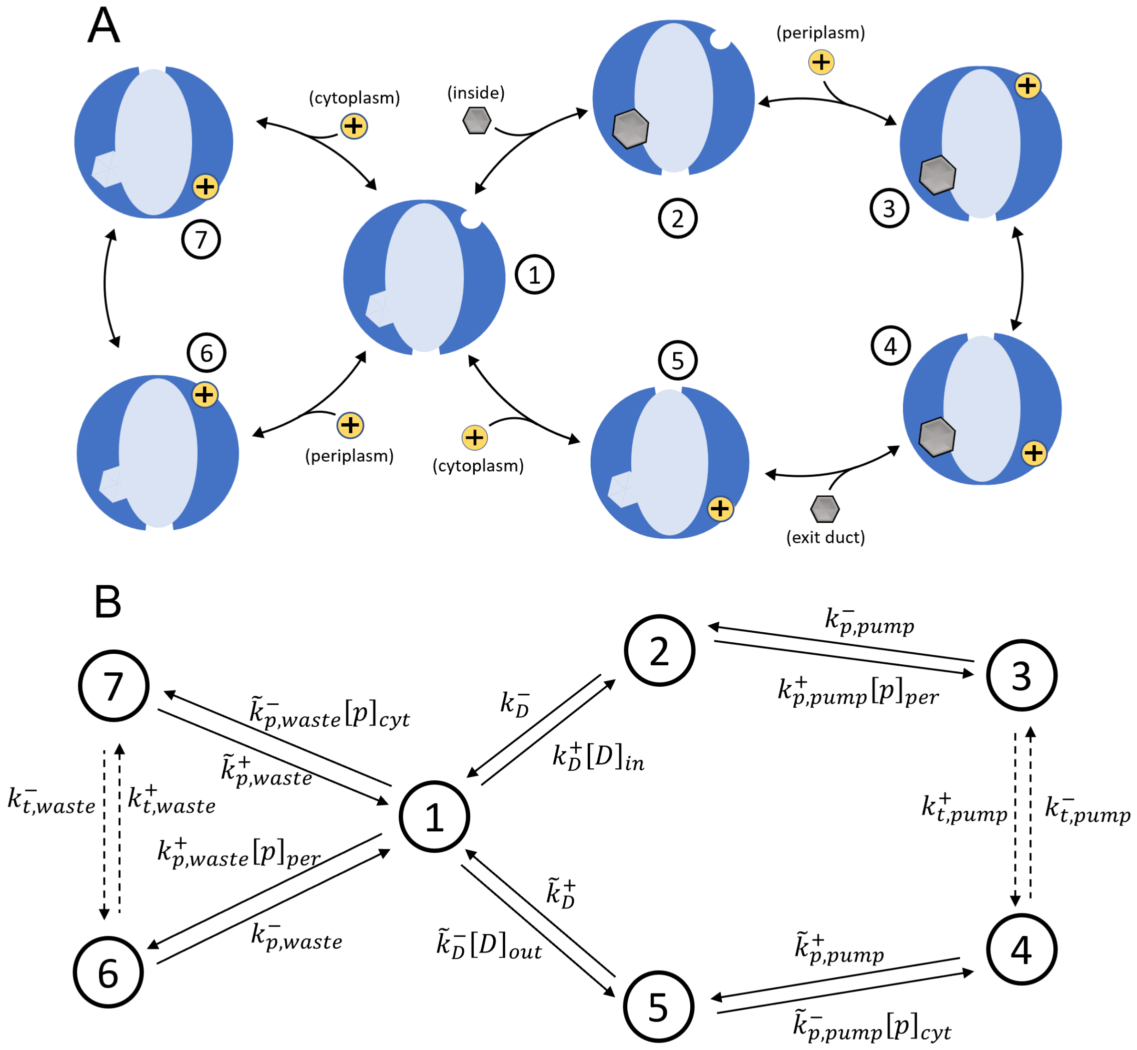


FIG. S3. **A.** Schematic diagram of the seven-state model for efflux pump operation, including a futile cycle by which one proton is transported into the cytoplasm without the associated extrusion of a drug molecule. **B.** The seven-state model represented as a network with each transition represented by a directed edge and labelled with its rate.

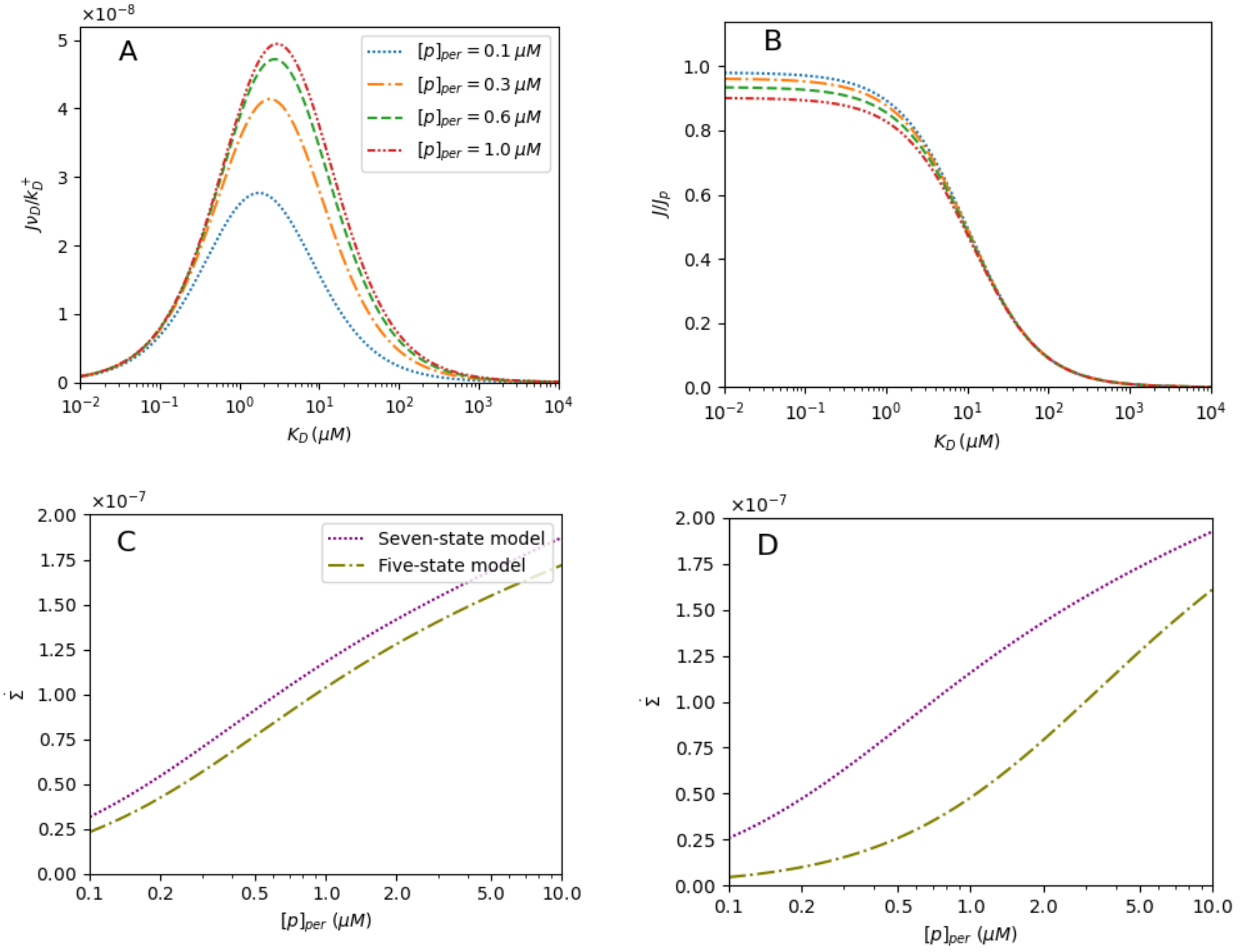


FIG. S4. **A.** The dimensionless efflux as a function of $K_D$, at varying periplasmic proton concentrations, for the seven-state model with a futile cycle. **B.** The chemical efficiency for the same model. **C.** The dimensionless entropy production rate for the seven- and five-state models as a function of the periplasmic proton concentration at $K_D = 10\ \mu M$. **D.** The analogous entropy production rate comparison at $K_D = 100\ \mu M$, probing the regime of weak binders in which drug efflux is suppressed. For the seven-state model, we set $K_{p,waste} = \tilde{K}_{p,waste} = K_{p,pump} = \tilde{K}_{p,pump} = 1\ \mu M$ in all panels, and we fix $\tilde{K}_D/K_D = 10$. Parameter values are otherwise the same as in Fig. S2.

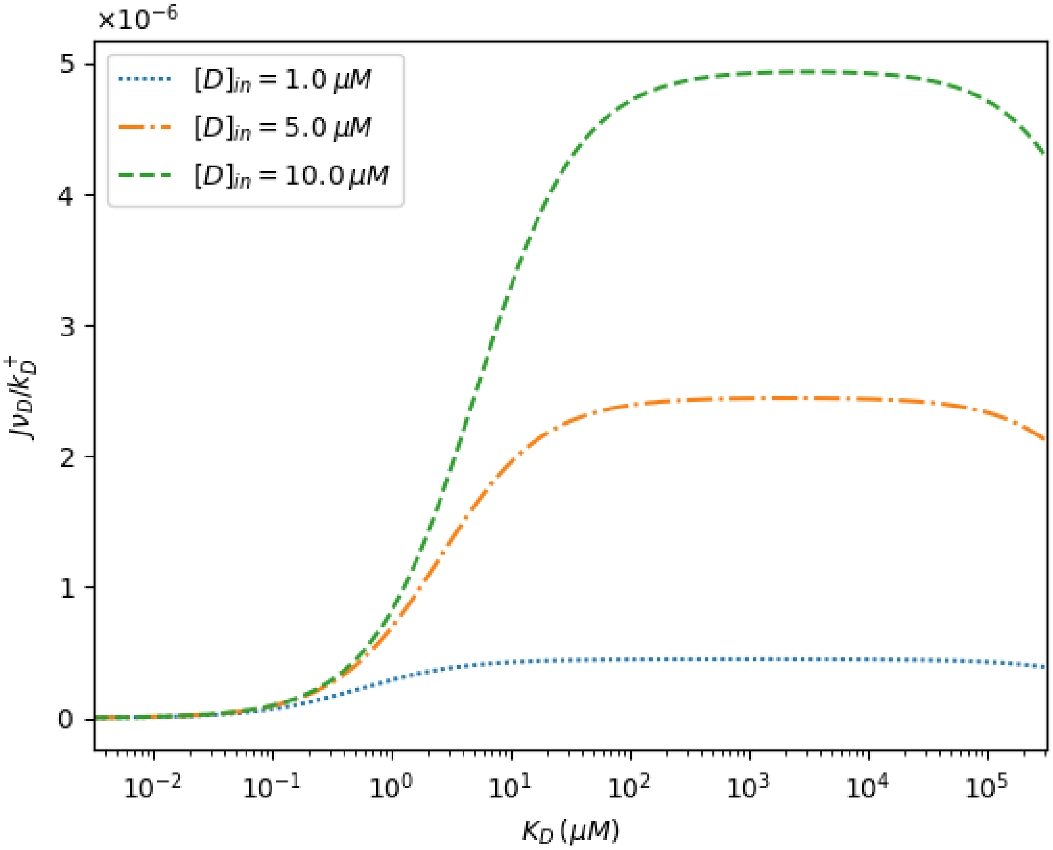


FIG. S5. Efflux as a function of $K_D$ at varying $[D]_{in}$ for a two-state, proton-independent passive channel as described in Sec. G. The distinct curves probe the behaviour at varying strengths of the drug concentration gradient, but show qualitatively similar behaviour. $[D]_{out} = 0.1\ \mu M$, $r_t^* = r_D = 10^8 s^{-1}$, and $\nu_D = 1 M^{-1}$.

[1] C. C. Su, H. Nikaido, and E. W. Yu, Ligand-transporter interaction in the AcrB multidrug efflux pump determined by fluorescence polarization assay, FEBS Letters **581**, 4972 (2007).

[2] I. Jarmoskaite, I. AlSadhan, P. P. Vaidyanathan, and D. Herschlag, How to measure and evaluate binding affinities, eLife **9**, e57264 (2020).

[3] T. Eicher, M. A. Seeger, C. Anselmi, W. Zhou, L. Brandstätter, F. Verrey, K. Diederichs, J. D. Faraldo-Gómez, and K. M. Pos, Coupling of remote alternating-access transport mechanisms for protons and substrates in the multidrug efflux pump AcrB, eLife **3**, 1 (2014).

[4] W. M. Haynes, ed., *CRC Handbook of Chemistry and Physics, 97th Ed.* (CRC Press, 2016).

[5] A. Panahi and C. L. Brooks, Membrane Environment Modulates the pKa Values of Transmembrane Helices, Journal of Physical Chemistry B **119**, 4601 (2015).

[6] D. Poland, Free energy of proton binding in proteins, Biopolymers **69**, 60 (2003).

[7] A. Fibich, K. Janko, and H.-J. Appel, Kinetics of proton binding to the sarcoplasmic reticulum ca-atpase in the e1 state, Biophys. J. **93**, 3092 (2007).

[8] A. H. Lang, C. K. Fisher, T. Mora, and P. Mehta, Thermodynamics of statistical inference by cells, Phys. Rev. Lett. **113**, 148103 (2014).

[9] J. D. Mallory, A. B. Kolomeisky, and O. A. Igoshin, Trade-offs between error, speed, noise, and energy dissipation in biological processes with proofreading, J. Phys. Chem. **123**, 4718 (2019).

[10] Q. Yu, J. D. Mallory, A. B. Kolomeisky, J. Ling, and O. A. Igoshin, Trade-offs between speed, accuracy, and dissipation in trnaile aminoacylation, J. Phys. Chem. Lett. **11**, 4001 (2020).
